# *SAMHD1* mutations in mantle cell lymphoma are recurrent and confer *in vitro* resistance to nucleoside analogues

**DOI:** 10.1101/2020.06.09.140376

**Authors:** Marco M Bühler, Junyan Lu, Sebastian Scheinost, Ferran Nadeu, Damien Roos-Weil, Manfred Hensel, Tharshika Thavayogarajah, Holger Moch, Markus G Manz, Eugenia Haralambieva, Ewerton Marques Maggio, Sílvia Beà, Eva Giné, Elías Campo, Olivier A. Bernard, Wolfgang Huber, Thorsten Zenz

## Abstract

The genomic landscape of mantle cell lymphoma (MCL) includes frequent alterations of *TP53*, *ATM*, *CCND1* and *KMT2D*. Thus far, the mutational landscape provides little information for treatment stratification. We characterized a cohort of MCL by targeted next generation sequencing and discovered *SAMHD1* as a novel recurrently mutated gene (8.5% of investigated cases, 4/47 samples). Furthermore, we provide evidence of in vitro resistance of *SAMHD1* mutated patient-derived MCL cells to cytarabine and fludarabine.

## Introduction

Mantle cell lymphoma (MCL) is an uncommon aggressive B-cell neoplasm which predominantly affects older adults and in the majority of cases is characterized by the t(11;14)(q13;q32) IGH*-CCND1* translocation, leading to cyclin D1 overexpression. Although *TP53* and *KMT2D* mutations provide prognostic information[1], mutations do not offer predictive value for treatment response in MCL (with the exception of *TP53*). To improve our knowledge of the mutational landscape in MCL, we used targeted next generation sequencing (NGS) and drug viability assays to explore drug susceptibility or resistance based on genomic alterations. We found *SAMHD1* (Sterile alpha motif and histidine aspartic domain containing deoxynucleoside triphosphate triphosphohydrolase 1) as a recurrently mutated gene (8.5% of investigated cases, 4/47 samples).

Furthermore, we provide evidence of *in vitro* resistance of *SAMHD1* mutated patient-derived MCL cells to cytarabine and fludarabine. *SAMHD1* is a gene which was first described to be mutated in Aicardi-Goutière Syndrome[2]. The known functions of *SAMHD1* include regulation of the intracellular dNTP pool and DNA repair[3]. Somatic *SAMHD1* mutations have been reported to occur in lymphoproliferative disorders such as chronic lymphocytic leukaemia (CLL), where mutations were found in 3% of treatment naïve and 11% of refractory/relapsed cases[4] and were detected at higher VAF after treatment[5]. Also, *SAMHD1* mutations were found in 18% of investigated cases of T-prolymphocytic leukaemia (T-PLL)[6].

## Methods

We sequenced a total of 47 MCL samples by means of targeted NGS (sample details listed in Supplementary Materials and Methods, covered regions listed in Supplementary Table 1). The study included peripheral blood samples from 25 MCL patients with leukemic disease and 22 frozen nodal or extranodal MCL tissue samples. Mutation calling was performed using a previously published pipeline[7]. Mean age of the cohort is 64.4 years (range 43-98) with a male predominance (83% of samples), 12 of the analysed samples represent pre-treated MCLs. Whole exome (WES) and whole genome sequencing (WGS) data was additionally available for 4 and 3 of samples, respectively. To explore the prognostic significance of *SAMHD1* mutations in MCL, samples with sufficient overall survival data from our cohort (n=35) were merged with cases from a recently published MCL study[8] (n=55), amounting to a total of 7 *SAMHD1* mutated and 83 wild type cases. To understand the impact of mutations (including *SAMHD1*) on the response to drugs, we exposed a set of patient-derived primary MCL cells (n=14, 11 *SAMHD1* wild type and 3 *SAMHD1* mutated) to increasing concentrations of cytarabine (ara-C), fludarabine, doxorubicine and nutlin-3a using the ATP-based CellTiter Glo assay (Promega), as previously described[9] (complete cell viability data provided in Supplementary Table 3). Associations between cell viability and genotype were identified by Student’s t-test (two-sided, equal variance) and p-values were adjusted for multiple testing using the Benjamini-Hochberg procedure. The study was approved by the Ethics Committee Heidelberg (University of Heidelberg, Germany; S-206/2011; S-356/2013) and the Cantonal Ethics Committee of Zurich (BASEC Nr. 2018-01618). Patients provided written informed consent prior to study.

## Results

In addition to mutations in known disease drivers such as *TP53* (n=19, 40.4%) and *ATM* (n=12, 25.5%), we identified mutations in *KMT2D* (n=6, 12.8%), *NFKBIE* (n=5, 10.6%) and *NOTCH1* (n=5, 10.6%) (Figure 1A and detailed list of all identified mutations provided in Supplementary Table 2). Additionally, we found *SAMHD1* mutations in four samples (n=4, 8.5%), two in previously treated and two in untreated patients. The mutational pattern is consistent with the hypothesized tumour suppressor function of *SAMHD1* with nonsense, missense and frameshift mutations (Figure 1B). All *SAMHD1* mutations had a high variant allele frequency (VAF, range 87-99%), indicating hemizygous or homozygous mutations. In three of the mutated samples, genomic data revealed deletion of chromosome 20q or uniparental disomy, explaining the high VAF. The presence of *SAMHD1* mutations was not associated with significant differences in overall survival (p=0.22, Figure 1C). Sample size of mutated cases was too small to explore associations between survival and different treatment protocols. Cell viability screening showed a significant difference in drug response to the nucleoside analogues cytarabine (adjusted p-value=1E-04) and fludarabine (adjusted p-value=4E-03), showing higher cell viability after 48 hours in *SAMHD1* mutated compared to unmutated patient-derived MCL cells (Figure 2A, Supplementary Tables 3-4 and Supplementary Figure 1), indicating that these mutations confer *in vitro* resistance. No significant differences in cell viability was observed in *TP53* mutated versus wild type samples for purine analogues, but *TP53* mutated MCL cells showed resistance for the *MDM2* inhibitor nutlin-3a (adjusted p-value 3E-03, Figure 2B and Supplementary Figure 1).

**Figure 1.**
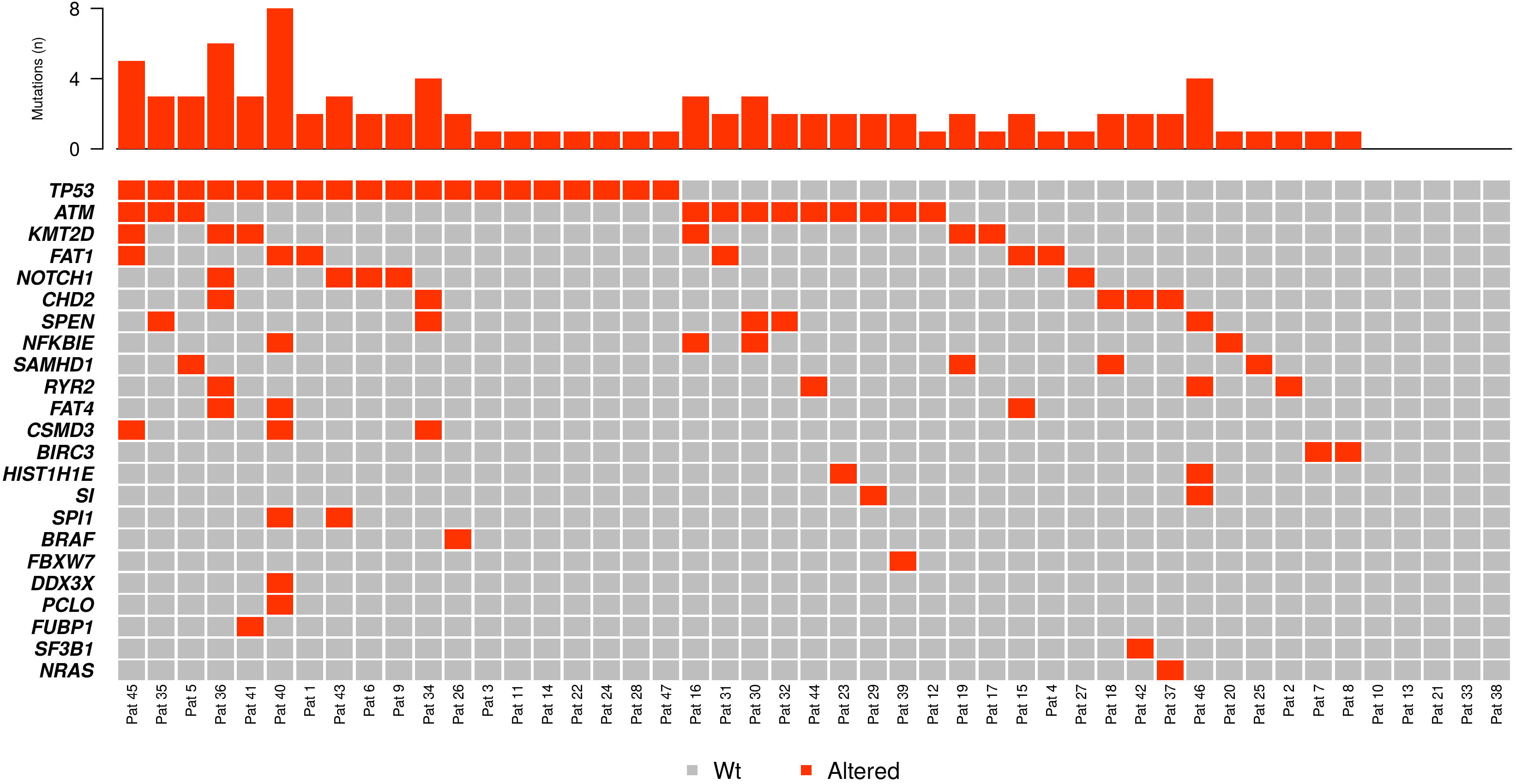
Mutations in the analyzed mantle cell lymphoma cohort and survival estimate. A) Gene mutations identified by targeted NGS sequencing (Oncoprint). Red cells represent mutations, top histogram indicates mutated genes per sample. B) Schematic representation of SAMHD1 with SAM (sterile alpha motif) and HD (histidine aspartic) domain. Symbols represent found SAMHD1 mutations: nonsense (red pentagon), missense (orange circles) and frameshift (purple square) mutations. C) Kaplan-Meier estimate of overall survival (OS) of 90 MCL patients, of which seven have mutated *SAMHD1* (blue line). Censored patients are indicated by vertical marks, no significant effect on survival was found (p=0.22).

**Figure 2.**
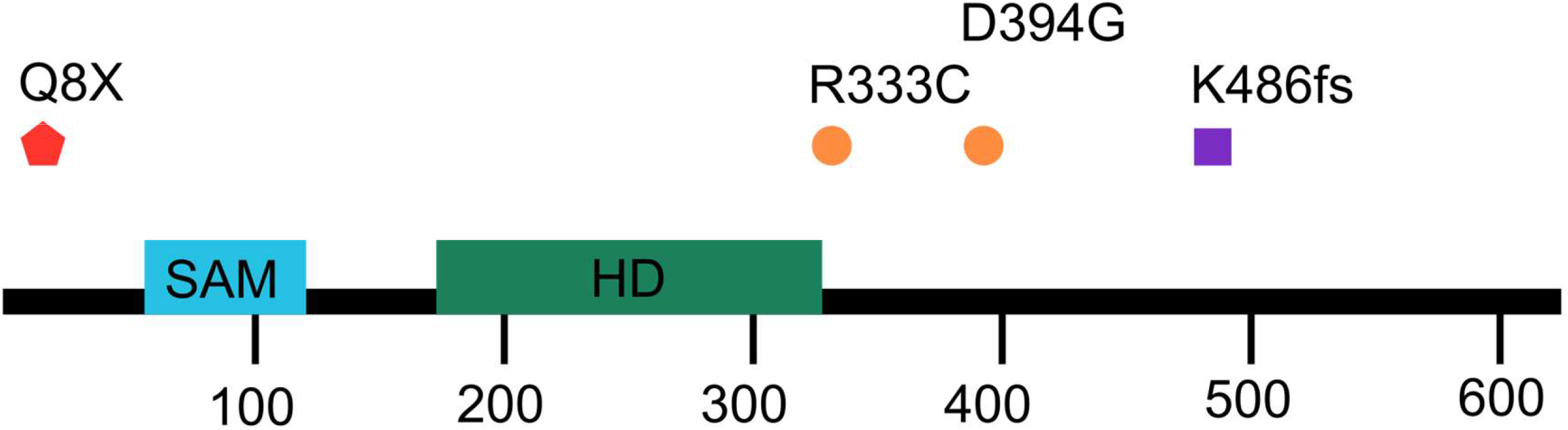
Effects of *SAMHD1* **and** *TP53* mutations on drug response. A) Boxplots illustrating cell viability after exposure to four different drugs (cytarabine, doxorubicine, fludarabine and nutlin-3a) by *SAMHD1* mutational status. Fraction of cells alive are shown (mean of 5 concentrations tested). *SAMHD1* mutations confer resistance to fludarabine (adjusted p-value 4E-04) and cytarabine (adjusted p-value 1E-04). B) Boxplots illustrating cell viability after exposure to four different drugs (cytarabine, doxorubicine, fludarabine and nutlin-3a) by *TP53* mutational status. Fraction of cells alive are shown (mean of 5 concentrations tested). *TP53* mutations confer resistance to nutlin-3a (adjusted p-value 3E-03).

## Discussion

In a recent MCL study[8], *SAMHD1* mutations were found in 5/82 cases (6.1% of samples) with two additional cases carrying *SAMHD1* deletions, furthermore Roider et al.[10] analysed the expression and mutational status of *SAMHD1* in patients of the MCL Younger and Elderly Trials and found mutations in 13/182 cases (7.1% of the samples), thus overall at a similar rate as in our study (8.5%) confirming that *SAMHD1* is recurrently mutated in MCL. Cytarabine is an efficient agent in the treatment of MCL[11], and our findings support further investigations on *SAMHD1* mutations as biomarkers of cytarabine resistance. For example, the reported higher incidence of *SAMHD1* mutations in refractory/relapsed CLL[4] could be due to selection of therapy resistant clones, since fludarabine is a commonly used agent (e.g. in the FCR regimen: fludarabine, cyclophosphamide, rituximab). We did not observe prognostic impact of *SAMHD1* mutations in our limited-sized cohort. Similarly neither SAMHD1 protein expression nor *SAMHD1* mutations were associated with complete remission (CR) or failure-free survival (FFS) rates in the aforementioned study of the MCL Younger and Elderly Trials[10], but higher SAMHD1 expression was associated with improved FFS in elderly patients treated with the R-CHOP (Rituximab, Cyclophosphamide, Doxorubicin, Vincristine, Prednisone) regimen. In acute myeloid leukaemia (AML) decreased SAMHD1 expression has been shown to be a biomarker that positively predicts response to cytarabine[12] and decitabine[13] treatment. Surprisingly, we found that *SAMHD1* mutations (which are predicted to impair its function) in MCL are associated with *in vitro* resistance to both cytarabine and fludarabine. Interestingly, Roider *et al*. provide evidence that high SAMHD1 expression in cell lines is associated with decreased susceptibility to Cytarabine exposure and that this effect was weakened when combining Cytarabine with Vincristine, hypothesizing that combination chemoimmunotherapy regimens (such as Hyper-CVAD) might overcome the cytarabine resistance in SAMHD1 mutated MCLs[10]. SAMHD1 is expressed at lower levels in B-cells compared to T-cells and myeloid cells[14]. A disruption by mutation or deletion could lead to an imbalance in nucleotide levels ultimately resulting in cytarabine resistance, for example by increasing dCTP levels, which compete with the metabolization of cytarabine to its active form ara-CTP, since elevated dNTP levels have been demonstrated in *SAMHD1* mutated T-PLL cells[6]. The observation that AML, T-PLL and MCL with altered *SAMHD1* expression (either by deletion, mutation or other mechanisms) respond differently might therefore be explained by differences in cell type and by different functional effect depending on type of alteration. A recent article by Davenne and co-workers[15] reported functional and pharmacological effects of SAMHD1 depletion in CLL cells, notably identifying forodesine as a potent agent in cells lacking SAMHD1 and elucidating the underlying mechanism. These results raise the possibility of a targeted use of forodesine in MCL carrying disrupting *SAMHD1* mutations, further highlighting the potential of SAMHD1 as a predictive biomarker. In conclusion, we report that *SAMHD1* is recurrently mutated in MCL and confers resistance to nucleoside analogue therapy *in vitro*. Further investigations are needed to potentially translate our *in vitro* result to clinical settings and to understand the resistance-conferring mechanism.

## Authorship

### Contribution

S.S. and D.R.W. performed the experiments; M.M.B., J.L., S.S., D.R.W., F.N., T.T., W.H., O.A.B. and T.Z. analysed the data; H.M., M.G.M., E.H., E.M.M., S.B., E.G., M.H. and E.C. contributed critical material; M.M.B. and T.Z. wrote the manuscript; T.Z. designed the study; All authors reviewed and edited the manuscript.

## Acknowledgements

M.M.B. is supported by a research grant of the Nuovo-Soldati Foundation for Cancer Research. T.Z. is supported by the Swiss Cancer League (KFS-4439-02-2018), the Dornonville de la Cour-Stiftung, the Cancer Research Center (CRC) of the Comprehensive Cancer Center Zurich (CCCZ) and the Clinical Research Priority Program (CRPP) of the University of Zurich (“Precision Hematology/Oncology”). O.A.B. acknowledges the support of JTC 2014-143 (INCA 2016-049).

## Data Sharing Statement

All identified mutations by targeted sequencing and cell viability data after drug exposure are listed in the Supplement. WES and methylation array data are available at the European Genome-phenome Archive under accession number EGAS00001001746.

---

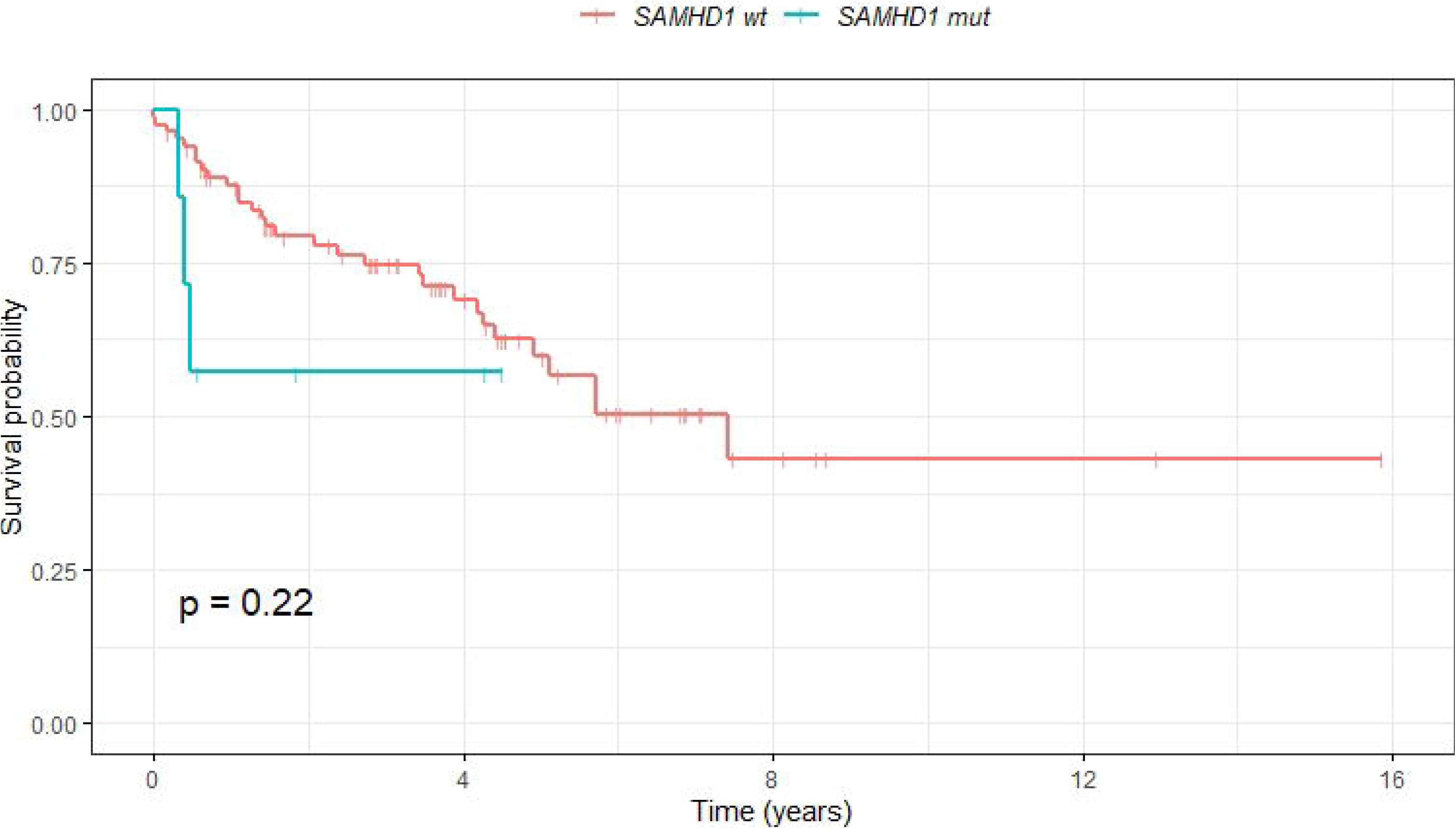

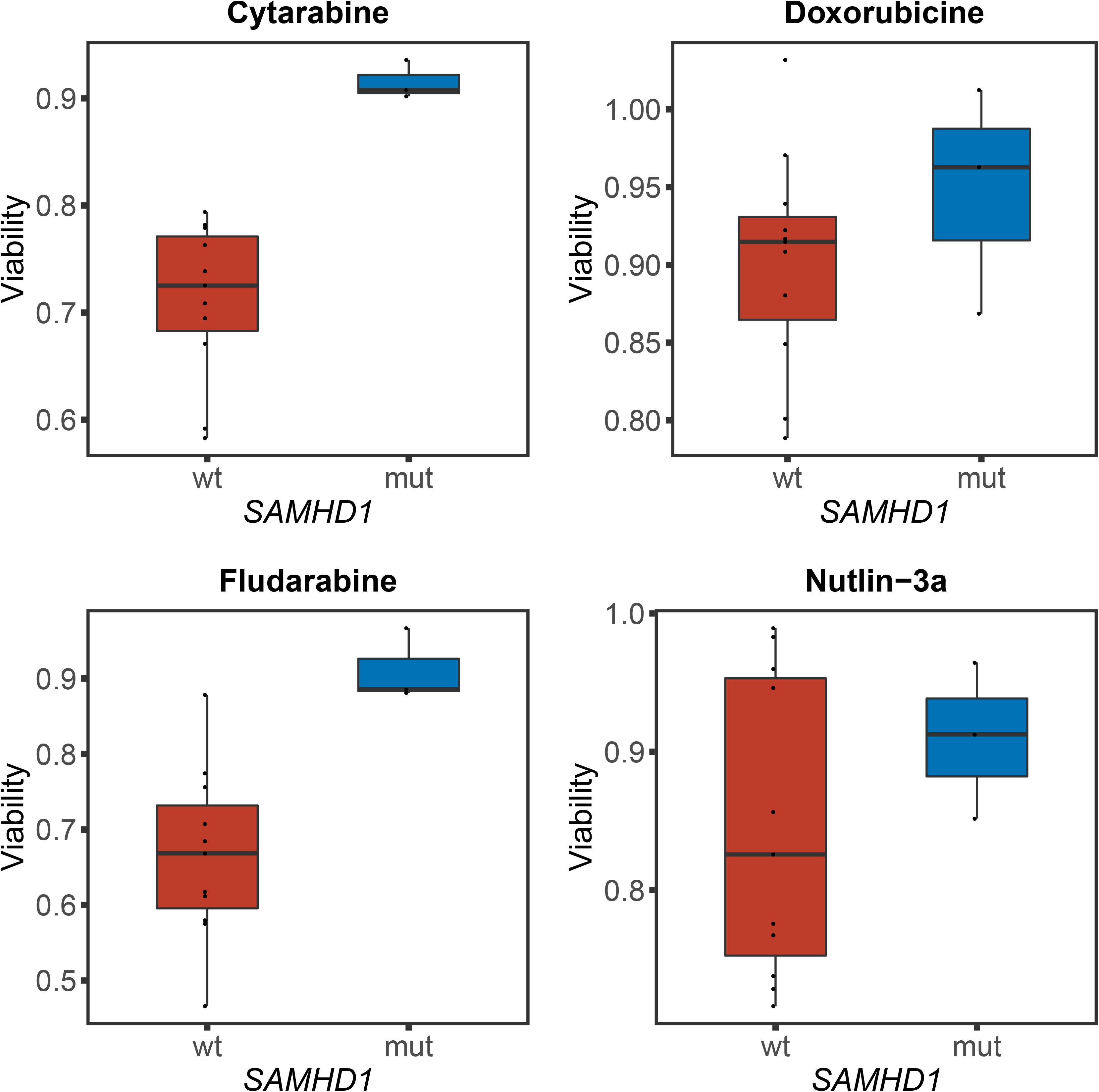

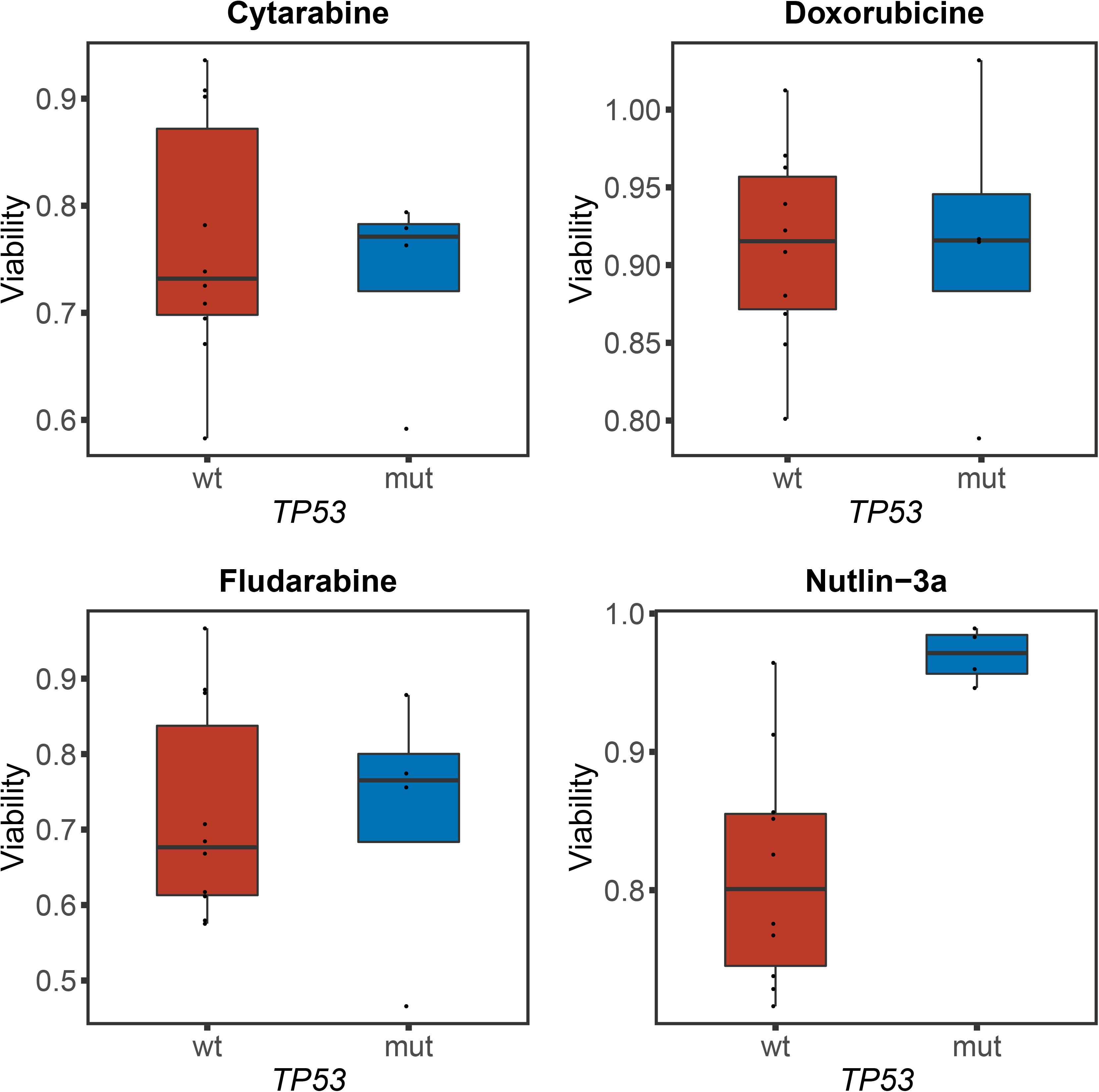

