## Supplemental Materials and Methods for "*SAMHD1* mutations in mantle cell lymphoma are recurrent and confer *in vitro* resistance to nucleoside analogues"

**Supplementary Materials and Methods**

**Patient samples**

We included peripheral blood samples from 25 mantle cell lymphoma (MCL) patients with leukemic disease. Blood was separated by a Ficoll gradient (GE Healthcare, Chicago, IL, USA), and mononuclear cells were cryopreserved. DNA was extracted using QIAmp DNA Kit (Qiagen, Germany) and DNA quantification was performed using a Qubit 2.0 Fluorometer (Life Technologies, Carlsbad, CA, USA). For drug response assays isolated mononuclear cells were washed in PBS (Invitrogen, Carlsbad, CA, USA), transferred into cryopreservation medium (55% RPMI 1640, 30% sterile-filtrated/heat inactivated fetal bovine serum (PAN Biotech, Germany) and 15% DMSO) and stored at -196°C. Additional fresh frozen tissue of 22 MCL nodal or extranodal tumor tissue samples diagnosed between 2001-2017 were obtained from the Biobank of the Department of Pathology and Molecular Pathology of the University Hospital of Zurich. DNA was extracted using the AllPrep DNA/RNA mini kit (Qiagen) and concentration was measured using a Nanodrop Spectrophotometer (ThermoFisher, Waltham, MA, USA). Mean age of the patient cohort (n=47) is 64.4 years (range 43-98) and 83% of patients are males. For association with survival, Kaplan-Meier estimates were calculated and plotted with R version 4.0 using the *survival* and *survminer* packages, log-rank test was used for comparison of survival between *SAMHD1* mutated and unmutated MCLs.

**Sequencing**

Targeted next generation sequencing was carried out on all 47 patients using two custom panels (sequenced regions see Supplementary Table 1). Ten ng of genomic DNA per primer pool were analyzed, libraries were generated with addition of NEXTflex paired-end adaptors (Bioo Scientific, Austin, TX, USA) before paired-end sequencing (2 x 250 bp reads) using an Illumina MiSeq flow cell and the onboard cluster method (Illumina, San Diego, CA, USA). Sequencing data was analyzed using a bioinformatic pipeline described in a previous study(1). Synonymous, intronic, UTR (excluding *NOTCH1* 3’UTR) variants as well as variants with a population frequency of >1% (according to 1000 Genomes project, ExAC and/or gnomAD) were filtered out. List of all found mutations are provided in Supplementary Table 2.

For 10 samples methylation array data was available. Whole exome sequencing (WES) and whole genome sequencing (WGS) data was available for 4 and 3 samples respectively. WES was carried out on a HiSeq 2000 instrument (Illumina) after library preparation on a SureSelect Automated Library Prep and Capture System (Agilent, Santa Clara, CA, USA)(2). WGS libraries were prepared according to manufacturer’s protocol using the TrueSeq Nano Library Preparation Kit (Illumina) and sequencing was carried out on a HiSeqX instrument (Illumina), as previously described(3,4).

**Drug response assay**

Drug response was assessed as previously described(2), briefly primary MCL cells of 14 patients (11 *SAMHD1* wildtype and 3 *SAMHD1* mutated and 10 *TP53* wildtype and 4 *TP53* mutated MCL) were exposed to four drugs: cytarabine (Sigma-Aldrich, St. Louis, MO, USA), fludarabine (Selleck Chemicals, Houston, TX, USA), doxorubicine (Selleck Chemicals) and nutlin-3a (Selleck Chemicals) in 5 concentrations each (five-fold serial dilution: 0.64 nM to 400 nM for doxorubicine and 16 nM to 10 µM for the other three drugs, respectively). After 48 hours, cell viability was assessed by luminescence measurement using the ATP-based CellTiter Glo assay (Promega, Madison, WI, USA) and the EnVision Multilabel Plate Reader (PerkinElmer, Waltham, MA, USA). Cell viability data is provided in Supplementary Table 3. Associations between cell viability and genotype were identified by Student’s t-test (two-sided, equal variance) and p-values were adjusted for multiple testing using the Benjamini-Hochberg procedure, significant results are summarized in Supplementary Table 4.
