## Supplementary Figure 1 for "*SAMHD1* mutations in mantle cell lymphoma are recurrent and confer *in vitro* resistance to nucleoside analogues"

Dose response plots showing cell viability at different concentrations, colors denote genotype (mutated in blue and wild type in red) of *SAMHD1* and *TP53* respectively.

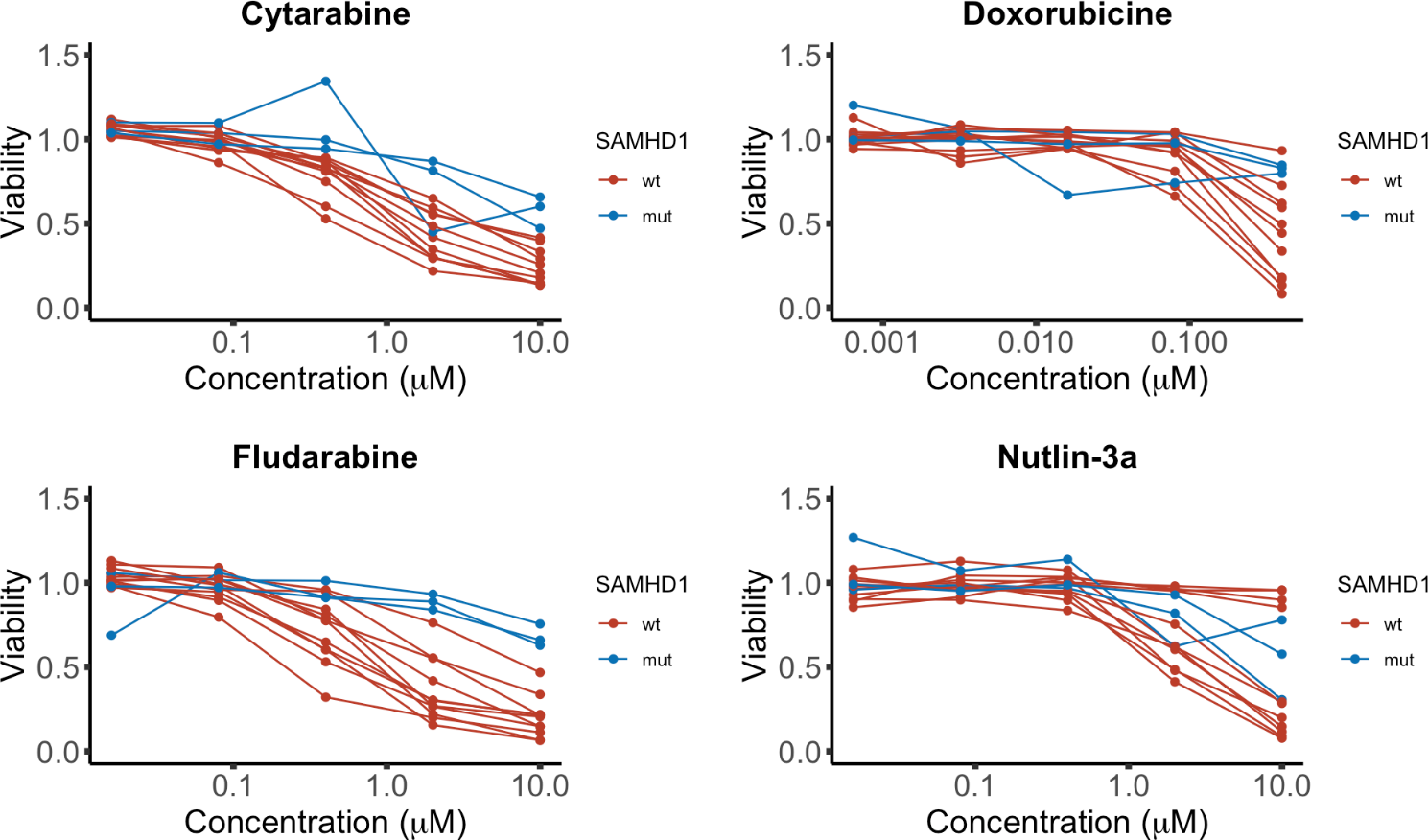

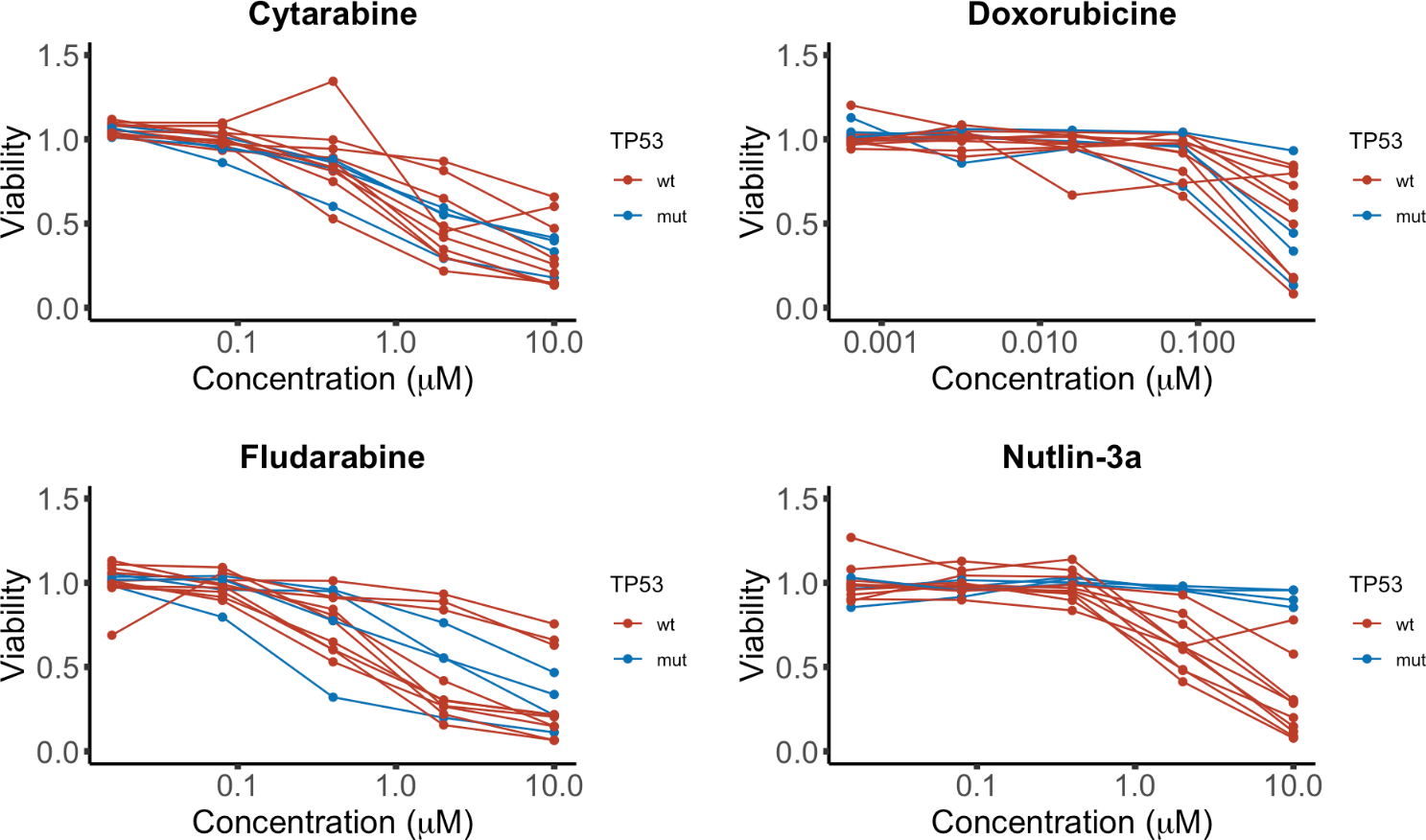
