## Supplementary Table 1 for "*SAMHD1* mutations in mantle cell lymphoma are recurrent and confer *in vitro* resistance to nucleoside analogues"

List of sequenced regions of the targeted next generation sequencing custom panels. Chromosome and base position are provided (hg19/GRCh37 genome assembly).

| chr1 | 16174516 | 16174640 | *SPEN* |  | chr9 | 37371639 | 37371859 | *PAX5* |
| --- | --- | --- | --- | --- | --- | --- | --- | --- |
| chr1 | 16199206 | 16199636 | *SPEN* |  | chr9 | 37371919 | 37372115 | *PAX5* |
| chr1 | 16202613 | 16203179 | *SPEN* |  | chr9 | 139390011 | 139390241 | *NOTCH1* |
| chr1 | 16235783 | 16236005 | *SPEN* |  | chr9 | 139390497 | 139390819 | *NOTCH1* |
| chr1 | 16237439 | 16237808 | *SPEN* |  | chr9 | 139390841 | 139391093 | *NOTCH1* |
| chr1 | 16242483 | 16242804 | *SPEN* |  | chr9 | 139391233 | 139392068 | *NOTCH1* |
| chr1 | 16245408 | 16245567 | *SPEN* |  | chr9 | 139393160 | 139393747 | *NOTCH1* |
| chr1 | 16245848 | 16246063 | *SPEN* |  | chr9 | 139394862 | 139395474 | *NOTCH1* |
| chr1 | 16247299 | 16247519 | *SPEN* |  | chr9 | 139396032 | 139396555 | *NOTCH1* |
| chr1 | 16248637 | 16248851 | *SPEN* |  | chr9 | 139396666 | 139396894 | *NOTCH1* |
| chr1 | 16254413 | 16258528 | *SPEN* |  | chr9 | 139396906 | 139397068 | *NOTCH1* |
| chr1 | 16258529 | 16262636 | *SPEN* |  | chr9 | 139397488 | 139397872 | *NOTCH1* |
| chr1 | 16262659 | 16262817 | *SPEN* |  | chr9 | 139399311 | 139399534 | *NOTCH1* |
| chr1 | 16263592 | 16264538 | *SPEN* |  | chr9 | 139399535 | 139399668 | *NOTCH1* |
| chr1 | 16265156 | 16265386 | *SPEN* |  | chr9 | 139399684 | 139400339 | *NOTCH1* |
| chr1 | 16265718 | 16265936 | *SPEN* |  | chr9 | 139400873 | 139401435 | *NOTCH1* |
| chr1 | 78414385 | 78414546 | *FUBP1* |  | chr9 | 139401646 | 139402024 | *NOTCH1* |
| chr1 | 78414799 | 78415015 | *FUBP1* |  | chr9 | 139402393 | 139402631 | *NOTCH1* |
| chr1 | 78420853 | 78421063 | *FUBP1* |  | chr9 | 139402647 | 139402930 | *NOTCH1* |
| chr1 | 78422230 | 78422412 | *FUBP1* |  | chr9 | 139403183 | 139403417 | *NOTCH1* |
| chr1 | 78425794 | 78426017 | *FUBP1* |  | chr9 | 139403426 | 139403585 | *NOTCH1* |
| chr1 | 78426024 | 78426232 | *FUBP1* |  | chr9 | 139404012 | 139404538 | *NOTCH1* |
| chr1 | 78428313 | 78428626 | *FUBP1* |  | chr9 | 139405072 | 139405331 | *NOTCH1* |
| chr1 | 78429208 | 78429428 | *FUBP1* |  | chr9 | 139405564 | 139405768 | *NOTCH1* |
| chr1 | 78429667 | 78430070 | *FUBP1* |  | chr9 | 139407335 | 139407689 | *NOTCH1* |
| chr1 | 78430319 | 78430480 | *FUBP1* |  | chr9 | 139407829 | 139408059 | *NOTCH1* |
| chr1 | 78430538 | 78430678 | *FUBP1* |  | chr9 | 139408933 | 139409140 | *NOTCH1* |
| chr1 | 78430718 | 78430943 | *FUBP1* |  | chr9 | 139409141 | 139409259 | *NOTCH1* |
| chr1 | 78432302 | 78432519 | *FUBP1* |  | chr9 | 139409609 | 139410190 | *NOTCH1* |
| chr1 | 78432546 | 78432869 | *FUBP1* |  | chr9 | 139410358 | 139410568 | *NOTCH1* |
| chr1 | 78433209 | 78433427 | *FUBP1* |  | chr9 | 139411749 | 139411897 | *NOTCH1* |
| chr1 | 78433706 | 78433916 | *FUBP1* |  | chr9 | 139412136 | 139412507 | *NOTCH1* |
| chr1 | 78435493 | 78435846 | *FUBP1* |  | chr9 | 139412526 | 139412918 | *NOTCH1* |
| chr1 | 78444502 | 78444726 | *FUBP1* |  | chr9 | 139412944 | 139413254 | *NOTCH1* |
| chr1 | 115256321 | 115256542 | *NRAS* |  | chr9 | 139413836 | 139414188 | *NOTCH1* |
| chr1 | 115258553 | 115258776 | *NRAS* |  | chr9 | 139417457 | 139417739 | *NOTCH1* |
| chr1 | 237205860 | 237206079 | *RYR2* |  | chr9 | 139418108 | 139418439 | *NOTCH1* |
| chr1 | 237433752 | 237433961 | *RYR2* |  | chr9 | 139438345 | 139438569 | *NOTCH1* |
| chr1 | 237494116 | 237494339 | *RYR2* |  | chr10 | 64572952 | 64574244 | *EGR2* |
| chr1 | 237519060 | 237519344 | *RYR2* |  | chr10 | 64573150 | 64573231 | *EGR2* |
| chr1 | 237527621 | 237527815 | *RYR2* |  | chr10 | 64573194 | 64573420 | *EGR2* |
| chr1 | 237532761 | 237532982 | *RYR2* |  | chr10 | 64573616 | 64573701 | *EGR2* |
| chr1 | 237537954 | 237538163 | *RYR2* |  | chr10 | 64573886 | 64574092 | *EGR2* |
| chr1 | 237540543 | 237540760 | *RYR2* |  | chr10 | 64575496 | 64575658 | *EGR2* |
| chr1 | 237550514 | 237550733 | *RYR2* |  | chr10 | 64575496 | 64575826 | *EGR2* |
| chr1 | 237551325 | 237551545 | *RYR2* |  | chr11 | 47376278 | 47376496 | *SPI1* |
| chr1 | 237580276 | 237580498 | *RYR2* |  | chr11 | 47376497 | 47376726 | *SPI1* |
| chr1 | 237586363 | 237586577 | *RYR2* |  | chr11 | 47376497 | 47376726 | *SPI1* |
| chr1 | 237604473 | 237604830 | *RYR2* |  | chr11 | 47376733 | 47377235 | *SPI1* |
| chr1 | 237608626 | 237608837 | *RYR2* |  | chr11 | 47376868 | 47377057 | *SPI1* |
| chr1 | 237617502 | 237617904 | *RYR2* |  | chr11 | 47380211 | 47380445 | *SPI1* |
| chr1 | 237619862 | 237620078 | *RYR2* |  | chr11 | 47380211 | 47380445 | *SPI1* |
| chr1 | 237632327 | 237632534 | *RYR2* |  | chr11 | 47380540 | 47380679 | *SPI1* |
| chr1 | 237655026 | 237655241 | *RYR2* |  | chr11 | 47381318 | 47381539 | *SPI1* |
| chr1 | 237656240 | 237656401 | *RYR2* |  | chr11 | 47381318 | 47381654 | *SPI1* |
| chr1 | 237659659 | 237660057 | *RYR2* |  | chr11 | 47397060 | 47397266 | *SPI1* |
| chr1 | 237663827 | 237664221 | *RYR2* |  | chr11 | 47397060 | 47397484 | *SPI1* |
| chr1 | 237666436 | 237666810 | *RYR2* |  | chr11 | 47399749 | 47399873 | *SPI1* |
| chr1 | 237669952 | 237670170 | *RYR2* |  | chr11 | 47399749 | 47400171 | *SPI1* |
| chr1 | 237674961 | 237675166 | *RYR2* |  | chr11 | 47400089 | 47400171 | *SPI1* |
| chr1 | 237693659 | 237693881 | *RYR2* |  | chr11 | 62559898 | 62560223 | *NXF1* |
| chr1 | 237711722 | 237711901 | *RYR2* |  | chr11 | 62560001 | 62560223 | *NXF1* |
| chr1 | 237713809 | 237714023 | *RYR2* |  | chr11 | 62561702 | 62561921 | *NXF1* |
| chr1 | 237729856 | 237730084 | *RYR2* |  | chr11 | 62562347 | 62562560 | *NXF1* |
| chr1 | 237732422 | 237732643 | *RYR2* |  | chr11 | 62562347 | 62562560 | *NXF1* |
| chr1 | 237752925 | 237753314 | *RYR2* |  | chr11 | 62563270 | 62563659 | *NXF1* |
| chr1 | 237753931 | 237754341 | *RYR2* |  | chr11 | 62563436 | 62563659 | *NXF1* |
| chr1 | 237754899 | 237755261 | *RYR2* |  | chr11 | 62563708 | 62564084 | *NXF1* |
| chr1 | 237756766 | 237756978 | *RYR2* |  | chr11 | 62563854 | 62564084 | *NXF1* |
| chr1 | 237758764 | 237758984 | *RYR2* |  | chr11 | 62564594 | 62564875 | *NXF1* |
| chr1 | 237765127 | 237765518 | *RYR2* |  | chr11 | 62564658 | 62564875 | *NXF1* |
| chr1 | 237773866 | 237774296 | *RYR2* |  | chr11 | 62565961 | 62566106 | *NXF1* |
| chr1 | 237777165 | 237778148 | *RYR2* |  | chr11 | 62567707 | 62567926 | *NXF1* |
| chr1 | 237780460 | 237780810 | *RYR2* |  | chr11 | 62567707 | 62568033 | *NXF1* |
| chr1 | 237787037 | 237787253 | *RYR2* |  | chr11 | 62568514 | 62568742 | *NXF1* |
| chr1 | 237788924 | 237789142 | *RYR2* |  | chr11 | 62568514 | 62569319 | *NXF1* |
| chr1 | 237790998 | 237791390 | *RYR2* |  | chr11 | 62568957 | 62569180 | *NXF1* |
| chr1 | 237794691 | 237794860 | *RYR2* |  | chr11 | 62569327 | 62569537 | *NXF1* |
| chr1 | 237796726 | 237797110 | *RYR2* |  | chr11 | 62569327 | 62569537 | *NXF1* |
| chr1 | 237798162 | 237798358 | *RYR2* |  | chr11 | 62569547 | 62569772 | *NXF1* |
| chr1 | 237801632 | 237801848 | *RYR2* |  | chr11 | 62570865 | 62571053 | *NXF1* |
| chr1 | 237802156 | 237802506 | *RYR2* |  | chr11 | 62570865 | 62571053 | *NXF1* |
| chr1 | 237804135 | 237804362 | *RYR2* |  | chr11 | 62571248 | 62571466 | *NXF1* |
| chr1 | 237806569 | 237806783 | *RYR2* |  | chr11 | 62572648 | 62572877 | *NXF1* |
| chr1 | 237811718 | 237811939 | *RYR2* |  | chr11 | 62572648 | 62572877 | *NXF1* |
| chr1 | 237813033 | 237813402 | *RYR2* |  | chr11 | 102195227 | 102196335 | *BIRC3* |
| chr1 | 237814703 | 237814899 | *RYR2* |  | chr11 | 102195432 | 102195592 | *BIRC3* |
| chr1 | 237817410 | 237817753 | *RYR2* |  | chr11 | 102195795 | 102196020 | *BIRC3* |
| chr1 | 237818986 | 237819353 | *RYR2* |  | chr11 | 102196170 | 102196335 | *BIRC3* |
| chr1 | 237821222 | 237821361 | *RYR2* |  | chr11 | 102198671 | 102198890 | *BIRC3* |
| chr1 | 237823275 | 237823484 | *RYR2* |  | chr11 | 102199442 | 102199650 | *BIRC3* |
| chr1 | 237824043 | 237824257 | *RYR2* |  | chr11 | 102199442 | 102199799 | *BIRC3* |
| chr1 | 237829736 | 237829956 | *RYR2* |  | chr11 | 102201562 | 102201760 | *BIRC3* |
| chr1 | 237831029 | 237831286 | *RYR2* |  | chr11 | 102201562 | 102202058 | *BIRC3* |
| chr1 | 237837324 | 237837539 | *RYR2* |  | chr11 | 102201843 | 102202058 | *BIRC3* |
| chr1 | 237837990 | 237838198 | *RYR2* |  | chr11 | 102206600 | 102207025 | *BIRC3* |
| chr1 | 237841263 | 237841479 | *RYR2* |  | chr11 | 102206809 | 102207025 | *BIRC3* |
| chr1 | 237843687 | 237843910 | *RYR2* |  | chr11 | 102207420 | 102208015 | *BIRC3* |
| chr1 | 237850645 | 237850872 | *RYR2* |  | chr11 | 102207609 | 102207814 | *BIRC3* |
| chr1 | 237862238 | 237862382 | *RYR2* |  | chr11 | 108098340 | 108098543 | *ATM* |
| chr1 | 237863374 | 237863805 | *RYR2* |  | chr11 | 108098340 | 108098712 | *ATM* |
| chr1 | 237865231 | 237865440 | *RYR2* |  | chr11 | 108099744 | 108099963 | *ATM* |
| chr1 | 237868498 | 237868656 | *RYR2* |  | chr11 | 108099744 | 108100070 | *ATM* |
| chr1 | 237870154 | 237870771 | *RYR2* |  | chr11 | 108106357 | 108106573 | *ATM* |
| chr1 | 237872005 | 237872415 | *RYR2* |  | chr11 | 108106357 | 108106573 | *ATM* |
| chr1 | 237872706 | 237872926 | *RYR2* |  | chr11 | 108114594 | 108114884 | *ATM* |
| chr1 | 237874996 | 237875213 | *RYR2* |  | chr11 | 108114708 | 108114884 | *ATM* |
| chr1 | 237880472 | 237880694 | *RYR2* |  | chr11 | 108115454 | 108115820 | *ATM* |
| chr1 | 237881722 | 237881935 | *RYR2* |  | chr11 | 108115601 | 108115820 | *ATM* |
| chr1 | 237886365 | 237886582 | *RYR2* |  | chr11 | 108117602 | 108117902 | *ATM* |
| chr1 | 237889468 | 237889689 | *RYR2* |  | chr11 | 108117812 | 108117902 | *ATM* |
| chr1 | 237890223 | 237890542 | *RYR2* |  | chr11 | 108119581 | 108119882 | *ATM* |
| chr1 | 237893549 | 237893675 | *RYR2* |  | chr11 | 108119700 | 108119882 | *ATM* |
| chr1 | 237895288 | 237895507 | *RYR2* |  | chr11 | 108121280 | 108121804 | *ATM* |
| chr1 | 237896942 | 237897150 | *RYR2* |  | chr11 | 108121484 | 108121593 | *ATM* |
| chr1 | 237905540 | 237905710 | *RYR2* |  | chr11 | 108122541 | 108122766 | *ATM* |
| chr1 | 237919517 | 237919742 | *RYR2* |  | chr11 | 108122541 | 108122766 | *ATM* |
| chr1 | 237920885 | 237921102 | *RYR2* |  | chr11 | 108123516 | 108123691 | *ATM* |
| chr1 | 237923001 | 237923211 | *RYR2* |  | chr11 | 108124510 | 108124634 | *ATM* |
| chr1 | 237924212 | 237924431 | *RYR2* |  | chr11 | 108124510 | 108124801 | *ATM* |
| chr1 | 237934037 | 237934258 | *RYR2* |  | chr11 | 108126803 | 108127019 | *ATM* |
| chr1 | 237935234 | 237935453 | *RYR2* |  | chr11 | 108126803 | 108127118 | *ATM* |
| chr1 | 237936757 | 237936977 | *RYR2* |  | chr11 | 108127992 | 108128214 | *ATM* |
| chr1 | 237941920 | 237942115 | *RYR2* |  | chr11 | 108127992 | 108128401 | *ATM* |
| chr1 | 237944786 | 237945001 | *RYR2* |  | chr11 | 108129702 | 108129875 | *ATM* |
| chr1 | 237946787 | 237948277 | *RYR2* |  | chr11 | 108129702 | 108129875 | *ATM* |
| chr1 | 237949202 | 237949419 | *RYR2* |  | chr11 | 108137892 | 108138108 | *ATM* |
| chr1 | 237951272 | 237951484 | *RYR2* |  | chr11 | 108139074 | 108139186 | *ATM* |
| chr1 | 237954685 | 237954873 | *RYR2* |  | chr11 | 108139074 | 108139350 | *ATM* |
| chr1 | 237955249 | 237955628 | *RYR2* |  | chr11 | 108141765 | 108141908 | *ATM* |
| chr1 | 237957124 | 237957340 | *RYR2* |  | chr11 | 108141765 | 108142148 | *ATM* |
| chr1 | 237958455 | 237958642 | *RYR2* |  | chr11 | 108142016 | 108142148 | *ATM* |
| chr1 | 237961317 | 237961531 | *RYR2* |  | chr11 | 108143213 | 108143458 | *ATM* |
| chr1 | 237965070 | 237965283 | *RYR2* |  | chr11 | 108143262 | 108143458 | *ATM* |
| chr1 | 237969378 | 237969595 | *RYR2* |  | chr11 | 108150160 | 108150358 | *ATM* |
| chr1 | 237972161 | 237972378 | *RYR2* |  | chr11 | 108151548 | 108151766 | *ATM* |
| chr1 | 237982312 | 237982516 | *RYR2* |  | chr11 | 108151548 | 108151912 | *ATM* |
| chr1 | 237991595 | 237991820 | *RYR2* |  | chr11 | 108153287 | 108153504 | *ATM* |
| chr1 | 237993723 | 237993951 | *RYR2* |  | chr11 | 108153287 | 108153681 | *ATM* |
| chr1 | 237994707 | 237994934 | *RYR2* |  | chr11 | 108154866 | 108154997 | *ATM* |
| chr1 | 237995788 | 237996009 | *RYR2* |  | chr11 | 108154866 | 108155307 | *ATM* |
| chr2 | 16083314 | 16083536 | *MYCN* |  | chr11 | 108155124 | 108155307 | *ATM* |
| chr2 | 16084469 | 16084696 | *MYCN* |  | chr11 | 108158277 | 108158492 | *ATM* |
| chr2 | 16086458 | 16086653 | *MYCN* |  | chr11 | 108159558 | 108159771 | *ATM* |
| chr2 | 47651179 | 47651365 | *MSH2* |  | chr11 | 108159558 | 108159974 | *ATM* |
| chr2 | 47656783 | 47657001 | *MSH2* |  | chr11 | 108160295 | 108160392 | *ATM* |
| chr2 | 47658339 | 47658557 | *MSH2* |  | chr11 | 108160295 | 108160552 | *ATM* |
| chr2 | 47663742 | 47663965 | *MSH2* |  | chr11 | 108163272 | 108163490 | *ATM* |
| chr2 | 47674821 | 47675045 | *MSH2* |  | chr11 | 108163272 | 108163669 | *ATM* |
| chr2 | 60713028 | 60713247 | *BCL11A* |  | chr11 | 108163918 | 108164134 | *ATM* |
| chr2 | 60714829 | 60715052 | *BCL11A* |  | chr11 | 108163918 | 108164308 | *ATM* |
| chr2 | 60715100 | 60715320 | *BCL11A* |  | chr11 | 108165624 | 108165830 | *ATM* |
| chr2 | 60723462 | 60723691 | *BCL11A* |  | chr11 | 108165624 | 108165830 | *ATM* |
| chr2 | 61119292 | 61119501 | *REL* |  | chr11 | 108167985 | 108168138 | *ATM* |
| chr2 | 61124838 | 61125037 | *REL* |  | chr11 | 108170335 | 108170501 | *ATM* |
| chr2 | 61129644 | 61129852 | *REL* |  | chr11 | 108170335 | 108170648 | *ATM* |
| chr2 | 61705921 | 61706142 | *XPO1* |  | chr11 | 108172332 | 108172549 | *ATM* |
| chr2 | 61708242 | 61708616 | *XPO1* |  | chr11 | 108172332 | 108172549 | *ATM* |
| chr2 | 61709496 | 61709863 | *XPO1* |  | chr11 | 108173383 | 108173791 | *ATM* |
| chr2 | 61710073 | 61710245 | *XPO1* |  | chr11 | 108173596 | 108173791 | *ATM* |
| chr2 | 61711049 | 61711263 | *XPO1* |  | chr11 | 108175393 | 108175606 | *ATM* |
| chr2 | 61712758 | 61713154 | *XPO1* |  | chr11 | 108178563 | 108178781 | *ATM* |
| chr2 | 61715267 | 61715439 | *XPO1* |  | chr11 | 108178563 | 108178781 | *ATM* |
| chr2 | 61715675 | 61715891 | *XPO1* |  | chr11 | 108180739 | 108181071 | *ATM* |
| chr2 | 61716428 | 61716646 | *XPO1* |  | chr11 | 108180930 | 108181071 | *ATM* |
| chr2 | 61717743 | 61717945 | *XPO1* |  | chr11 | 108183107 | 108183279 | *ATM* |
| chr2 | 61719091 | 61719667 | *XPO1* |  | chr11 | 108186544 | 108186671 | *ATM* |
| chr2 | 61719699 | 61719892 | *XPO1* |  | chr11 | 108186544 | 108186671 | *ATM* |
| chr2 | 61719913 | 61720295 | *XPO1* |  | chr11 | 108186702 | 108186876 | *ATM* |
| chr2 | 61720844 | 61721262 | *XPO1* |  | chr11 | 108187983 | 108188150 | *ATM* |
| chr2 | 61722554 | 61722761 | *XPO1* |  | chr11 | 108187983 | 108188274 | *ATM* |
| chr2 | 61723973 | 61724163 | *XPO1* |  | chr11 | 108190628 | 108190830 | *ATM* |
| chr2 | 61725784 | 61726128 | *XPO1* |  | chr11 | 108190628 | 108190830 | *ATM* |
| chr2 | 61726820 | 61727035 | *XPO1* |  | chr11 | 108191960 | 108192178 | *ATM* |
| chr2 | 61729083 | 61729269 | *XPO1* |  | chr11 | 108195931 | 108196113 | *ATM* |
| chr2 | 61729305 | 61729523 | *XPO1* |  | chr11 | 108195931 | 108196285 | *ATM* |
| chr2 | 61735400 | 61735625 | *XPO1* |  | chr11 | 108196699 | 108196921 | *ATM* |
| chr2 | 61749637 | 61749858 | *XPO1* |  | chr11 | 108196699 | 108197133 | *ATM* |
| chr2 | 61753513 | 61753709 | *XPO1* |  | chr11 | 108198324 | 108198521 | *ATM* |
| chr2 | 61756885 | 61757108 | *XPO1* |  | chr11 | 108198324 | 108198521 | *ATM* |
| chr2 | 61760872 | 61761061 | *XPO1* |  | chr11 | 108199621 | 108199989 | *ATM* |
| chr2 | 61761901 | 61762128 | *XPO1* |  | chr11 | 108199773 | 108199989 | *ATM* |
| chr2 | 61765023 | 61765213 | *XPO1* |  | chr11 | 108200768 | 108201153 | *ATM* |
| chr2 | 140990600 | 140990928 | *LRP1B* |  | chr11 | 108200983 | 108201153 | *ATM* |
| chr2 | 140992321 | 140992539 | *LRP1B* |  | chr11 | 108202020 | 108202426 | *ATM* |
| chr2 | 140995545 | 140995972 | *LRP1B* |  | chr11 | 108202232 | 108202426 | *ATM* |
| chr2 | 140996991 | 140997205 | *LRP1B* |  | chr11 | 108202469 | 108202781 | *ATM* |
| chr2 | 141004584 | 141004800 | *LRP1B* |  | chr11 | 108202665 | 108202781 | *ATM* |
| chr2 | 141027755 | 141027971 | *LRP1B* |  | chr11 | 108203456 | 108203658 | *ATM* |
| chr2 | 141031953 | 141032325 | *LRP1B* |  | chr11 | 108204554 | 108204773 | *ATM* |
| chr2 | 141055155 | 141055596 | *LRP1B* |  | chr11 | 108204554 | 108204773 | *ATM* |
| chr2 | 141072324 | 141072680 | *LRP1B* |  | chr11 | 108205578 | 108205843 | *ATM* |
| chr2 | 141079344 | 141079729 | *LRP1B* |  | chr11 | 108205722 | 108205843 | *ATM* |
| chr2 | 141081333 | 141081679 | *LRP1B* |  | chr11 | 108206502 | 108206732 | *ATM* |
| chr2 | 141083124 | 141083452 | *LRP1B* |  | chr11 | 108213801 | 108214008 | *ATM* |
| chr2 | 141091971 | 141092172 | *LRP1B* |  | chr11 | 108213801 | 108214135 | *ATM* |
| chr2 | 141092983 | 141093412 | *LRP1B* |  | chr11 | 108216335 | 108216534 | *ATM* |
| chr2 | 141108202 | 141108804 | *LRP1B* |  | chr11 | 108216335 | 108216650 | *ATM* |
| chr2 | 141110403 | 141110747 | *LRP1B* |  | chr11 | 108217945 | 108218145 | *ATM* |
| chr2 | 141113729 | 141114146 | *LRP1B* |  | chr11 | 108217945 | 108218145 | *ATM* |
| chr2 | 141115511 | 141115727 | *LRP1B* |  | chr11 | 108224422 | 108224646 | *ATM* |
| chr2 | 141116348 | 141116549 | *LRP1B* |  | chr11 | 108225461 | 108225678 | *ATM* |
| chr2 | 141122178 | 141122401 | *LRP1B* |  | chr11 | 108225461 | 108225678 | *ATM* |
| chr2 | 141128239 | 141128453 | *LRP1B* |  | chr11 | 108235743 | 108235962 | *ATM* |
| chr2 | 141128622 | 141128838 | *LRP1B* |  | chr11 | 108236025 | 108236246 | *ATM* |
| chr2 | 141128854 | 141129073 | *LRP1B* |  | chr11 | 108236025 | 108236246 | *ATM* |
| chr2 | 141130422 | 141130745 | *LRP1B* |  | chr12 | 25398183 | 25398405 | *KRAS* |
| chr2 | 141135725 | 141135949 | *LRP1B* |  | chr12 | 25398183 | 25398405 | *KRAS* |
| chr2 | 141143409 | 141143619 | *LRP1B* |  | chr12 | 49415518 | 49415739 | *KMT2D* |
| chr2 | 141199988 | 141200206 | *LRP1B* |  | chr12 | 49415757 | 49415980 | *KMT2D* |
| chr2 | 141201882 | 141202050 | *LRP1B* |  | chr12 | 49415757 | 49416200 | *KMT2D* |
| chr2 | 141202063 | 141202273 | *LRP1B* |  | chr12 | 49416250 | 49416452 | *KMT2D* |
| chr2 | 141208072 | 141208293 | *LRP1B* |  | chr12 | 49416250 | 49416663 | *KMT2D* |
| chr2 | 141213853 | 141214228 | *LRP1B* |  | chr12 | 49418310 | 49418535 | *KMT2D* |
| chr2 | 141214871 | 141215227 | *LRP1B* |  | chr12 | 49418310 | 49418535 | *KMT2D* |
| chr2 | 141232713 | 141232917 | *LRP1B* |  | chr12 | 49418556 | 49418772 | *KMT2D* |
| chr2 | 141242812 | 141243118 | *LRP1B* |  | chr12 | 49419901 | 49420081 | *KMT2D* |
| chr2 | 141245087 | 141245361 | *LRP1B* |  | chr12 | 49419901 | 49421139 | *KMT2D* |
| chr2 | 141250108 | 141250327 | *LRP1B* |  | chr12 | 49420285 | 49420502 | *KMT2D* |
| chr2 | 141252992 | 141253412 | *LRP1B* |  | chr12 | 49420652 | 49420794 | *KMT2D* |
| chr2 | 141259138 | 141259453 | *LRP1B* |  | chr12 | 49420842 | 49421064 | *KMT2D* |
| chr2 | 141260344 | 141260715 | *LRP1B* |  | chr12 | 49421490 | 49421720 | *KMT2D* |
| chr2 | 141264296 | 141264512 | *LRP1B* |  | chr12 | 49421490 | 49421720 | *KMT2D* |
| chr2 | 141267458 | 141267665 | *LRP1B* |  | chr12 | 49421736 | 49421964 | *KMT2D* |
| chr2 | 141272216 | 141272366 | *LRP1B* |  | chr12 | 49422529 | 49422755 | *KMT2D* |
| chr2 | 141274389 | 141274732 | *LRP1B* |  | chr12 | 49422529 | 49423024 | *KMT2D* |
| chr2 | 141283397 | 141283610 | *LRP1B* |  | chr12 | 49422936 | 49423024 | *KMT2D* |
| chr2 | 141283742 | 141283957 | *LRP1B* |  | chr12 | 49423119 | 49423327 | *KMT2D* |
| chr2 | 141291525 | 141291718 | *LRP1B* |  | chr12 | 49424031 | 49424255 | *KMT2D* |
| chr2 | 141294099 | 141294297 | *LRP1B* |  | chr12 | 49424031 | 49424255 | *KMT2D* |
| chr2 | 141298363 | 141298790 | *LRP1B* |  | chr12 | 49424351 | 49424572 | *KMT2D* |
| chr2 | 141299184 | 141299581 | *LRP1B* |  | chr12 | 49424594 | 49424822 | *KMT2D* |
| chr2 | 141356098 | 141356459 | *LRP1B* |  | chr12 | 49424594 | 49424822 | *KMT2D* |
| chr2 | 141358829 | 141359252 | *LRP1B* |  | chr12 | 49424882 | 49425512 | *KMT2D* |
| chr2 | 141457651 | 141458195 | *LRP1B* |  | chr12 | 49425079 | 49425273 | *KMT2D* |
| chr2 | 141459107 | 141459422 | *LRP1B* |  | chr12 | 49425350 | 49425512 | *KMT2D* |
| chr2 | 141459616 | 141460156 | *LRP1B* |  | chr12 | 49425547 | 49425837 | *KMT2D* |
| chr2 | 141473337 | 141473564 | *LRP1B* |  | chr12 | 49425606 | 49425837 | *KMT2D* |
| chr2 | 141473565 | 141473793 | *LRP1B* |  | chr12 | 49425852 | 49427177 | *KMT2D* |
| chr2 | 141474229 | 141474416 | *LRP1B* |  | chr12 | 49425922 | 49426144 | *KMT2D* |
| chr2 | 141526690 | 141527047 | *LRP1B* |  | chr12 | 49426311 | 49426406 | *KMT2D* |
| chr2 | 141528387 | 141528659 | *LRP1B* |  | chr12 | 49426537 | 49426622 | *KMT2D* |
| chr2 | 141533499 | 141533907 | *LRP1B* |  | chr12 | 49426809 | 49426993 | *KMT2D* |
| chr2 | 141571207 | 141571408 | *LRP1B* |  | chr12 | 49427206 | 49427365 | *KMT2D* |
| chr2 | 141597414 | 141597768 | *LRP1B* |  | chr12 | 49427206 | 49428505 | *KMT2D* |
| chr2 | 141598354 | 141598689 | *LRP1B* |  | chr12 | 49427506 | 49427727 | *KMT2D* |
| chr2 | 141607494 | 141607913 | *LRP1B* |  | chr12 | 49427845 | 49428072 | *KMT2D* |
| chr2 | 141609097 | 141609373 | *LRP1B* |  | chr12 | 49428124 | 49428328 | *KMT2D* |
| chr2 | 141625038 | 141625425 | *LRP1B* |  | chr12 | 49428566 | 49428756 | *KMT2D* |
| chr2 | 141625481 | 141625886 | *LRP1B* |  | chr12 | 49428566 | 49428756 | *KMT2D* |
| chr2 | 141641196 | 141641595 | *LRP1B* |  | chr12 | 49430855 | 49431725 | *KMT2D* |
| chr2 | 141643564 | 141643937 | *LRP1B* |  | chr12 | 49431074 | 49431274 | *KMT2D* |
| chr2 | 141660436 | 141660759 | *LRP1B* |  | chr12 | 49431395 | 49431496 | *KMT2D* |
| chr2 | 141665242 | 141665663 | *LRP1B* |  | chr12 | 49431625 | 49431725 | *KMT2D* |
| chr2 | 141680508 | 141680724 | *LRP1B* |  | chr12 | 49431747 | 49432784 | *KMT2D* |
| chr2 | 141707668 | 141707976 | *LRP1B* |  | chr12 | 49431922 | 49432136 | *KMT2D* |
| chr2 | 141709364 | 141709574 | *LRP1B* |  | chr12 | 49432311 | 49432501 | *KMT2D* |
| chr2 | 141739669 | 141739875 | *LRP1B* |  | chr12 | 49432625 | 49432784 | *KMT2D* |
| chr2 | 141746899 | 141747331 | *LRP1B* |  | chr12 | 49432946 | 49433169 | *KMT2D* |
| chr2 | 141751517 | 141751737 | *LRP1B* |  | chr12 | 49433188 | 49433411 | *KMT2D* |
| chr2 | 141762864 | 141763060 | *LRP1B* |  | chr12 | 49433188 | 49433411 | *KMT2D* |
| chr2 | 141770944 | 141771324 | *LRP1B* |  | chr12 | 49433438 | 49434100 | *KMT2D* |
| chr2 | 141773115 | 141773524 | *LRP1B* |  | chr12 | 49433606 | 49433817 | *KMT2D* |
| chr2 | 141777293 | 141777853 | *LRP1B* |  | chr12 | 49433874 | 49434100 | *KMT2D* |
| chr2 | 141806462 | 141806796 | *LRP1B* |  | chr12 | 49434130 | 49434815 | *KMT2D* |
| chr2 | 141812670 | 141812843 | *LRP1B* |  | chr12 | 49434213 | 49434444 | *KMT2D* |
| chr2 | 141816445 | 141816658 | *LRP1B* |  | chr12 | 49434599 | 49434815 | *KMT2D* |
| chr2 | 141819449 | 141819848 | *LRP1B* |  | chr12 | 49434832 | 49435383 | *KMT2D* |
| chr2 | 141945834 | 141946158 | *LRP1B* |  | chr12 | 49435046 | 49435168 | *KMT2D* |
| chr2 | 141986642 | 141987162 | *LRP1B* |  | chr12 | 49435406 | 49435543 | *KMT2D* |
| chr2 | 142004726 | 142004947 | *LRP1B* |  | chr12 | 49435406 | 49435543 | *KMT2D* |
| chr2 | 142011893 | 142012303 | *LRP1B* |  | chr12 | 49435636 | 49436165 | *KMT2D* |
| chr2 | 142237959 | 142238156 | *LRP1B* |  | chr12 | 49435748 | 49435971 | *KMT2D* |
| chr2 | 142567802 | 142568020 | *LRP1B* |  | chr12 | 49436291 | 49436481 | *KMT2D* |
| chr2 | 142888135 | 142888403 | *LRP1B* |  | chr12 | 49436291 | 49436690 | *KMT2D* |
| chr2 | 150000405 | 150000622 | *LYPD6B* |  | chr12 | 49436834 | 49437058 | *KMT2D* |
| chr2 | 150000698 | 150000860 | *LYPD6B* |  | chr12 | 49436834 | 49437330 | *KMT2D* |
| chr2 | 150001723 | 150001948 | *LYPD6B* |  | chr12 | 49437194 | 49437330 | *KMT2D* |
| chr2 | 150002010 | 150002240 | *LYPD6B* |  | chr12 | 49437368 | 49437795 | *KMT2D* |
| chr2 | 150002337 | 150002560 | *LYPD6B* |  | chr12 | 49437566 | 49437795 | *KMT2D* |
| chr2 | 150002622 | 150002841 | *LYPD6B* |  | chr12 | 49437877 | 49438319 | *KMT2D* |
| chr2 | 150029600 | 150029817 | *LYPD6B* |  | chr12 | 49438093 | 49438319 | *KMT2D* |
| chr2 | 150057699 | 150057923 | *LYPD6B* |  | chr12 | 49438338 | 49438753 | *KMT2D* |
| chr2 | 150086470 | 150086612 | *-* |  | chr12 | 49438539 | 49438753 | *KMT2D* |
| chr2 | 150098456 | 150098665 | *-* |  | chr12 | 49439612 | 49439979 | *KMT2D* |
| chr2 | 198256922 | 198257304 | *SF3B1* |  | chr12 | 49439770 | 49439979 | *KMT2D* |
| chr2 | 198257492 | 198257709 | *SF3B1* |  | chr12 | 49440024 | 49440226 | *KMT2D* |
| chr2 | 198257714 | 198257927 | *SF3B1* |  | chr12 | 49440386 | 49440579 | *KMT2D* |
| chr2 | 198260669 | 198261057 | *SF3B1* |  | chr12 | 49440386 | 49440579 | *KMT2D* |
| chr2 | 198262577 | 198262949 | *SF3B1* |  | chr12 | 49441691 | 49441910 | *KMT2D* |
| chr2 | 198263137 | 198263351 | *SF3B1* |  | chr12 | 49442378 | 49442602 | *KMT2D* |
| chr2 | 198264769 | 198265179 | *SF3B1* |  | chr12 | 49442378 | 49442602 | *KMT2D* |
| chr2 | 198265266 | 198265680 | *SF3B1* |  | chr12 | 49442848 | 49443043 | *KMT2D* |
| chr2 | 198266076 | 198266294 | *SF3B1* |  | chr12 | 49443369 | 49443579 | *KMT2D* |
| chr2 | 198266322 | 198266859 | *SF3B1* |  | chr12 | 49443369 | 49444588 | *KMT2D* |
| chr2 | 198267150 | 198267555 | *SF3B1* |  | chr12 | 49443752 | 49443979 | *KMT2D* |
| chr2 | 198267585 | 198267796 | *SF3B1* |  | chr12 | 49444127 | 49444317 | *KMT2D* |
| chr2 | 198268150 | 198268493 | *SF3B1* |  | chr12 | 49444413 | 49444588 | *KMT2D* |
| chr2 | 198269793 | 198270028 | *SF3B1* |  | chr12 | 49444618 | 49445176 | *KMT2D* |
| chr2 | 198270073 | 198270292 | *SF3B1* |  | chr12 | 49444694 | 49444913 | *KMT2D* |
| chr2 | 198272618 | 198273417 | *SF3B1* |  | chr12 | 49445091 | 49445176 | *KMT2D* |
| chr2 | 198274405 | 198274744 | *SF3B1* |  | chr12 | 49445335 | 49446212 | *KMT2D* |
| chr2 | 198281414 | 198281667 | *SF3B1* |  | chr12 | 49445553 | 49445775 | *KMT2D* |
| chr2 | 198283180 | 198283365 | *SF3B1* |  | chr12 | 49445943 | 49446044 | *KMT2D* |
| chr2 | 198283521 | 198283742 | *SF3B1* |  | chr12 | 49446303 | 49446529 | *KMT2D* |
| chr2 | 198284965 | 198285327 | *SF3B1* |  | chr12 | 49446303 | 49446529 | *KMT2D* |
| chr2 | 198285614 | 198285944 | *SF3B1* |  | chr12 | 49446664 | 49446888 | *KMT2D* |
| chr2 | 198288369 | 198288720 | *SF3B1* |  | chr12 | 49446940 | 49447157 | *KMT2D* |
| chr2 | 198299634 | 198299857 | *SF3B1* |  | chr12 | 49446940 | 49447157 | *KMT2D* |
| chr3 | 38180102 | 38180585 | *MYD88* |  | chr12 | 49447231 | 49447452 | *KMT2D* |
| chr3 | 38181308 | 38181536 | *MYD88* |  | chr12 | 49447741 | 49447967 | *KMT2D* |
| chr3 | 38181736 | 38182172 | *MYD88* |  | chr12 | 49447741 | 49447967 | *KMT2D* |
| chr3 | 38182217 | 38182420 | *MYD88* |  | chr12 | 49448019 | 49448540 | *KMT2D* |
| chr3 | 38182459 | 38182799 | *MYD88* |  | chr12 | 49448117 | 49448345 | *KMT2D* |
| chr3 | 164697145 | 164697345 | *SI* |  | chr12 | 49448634 | 49448856 | *KMT2D* |
| chr3 | 164699921 | 164700214 | *SI* |  | chr12 | 49448634 | 49448856 | *KMT2D* |
| chr3 | 164700771 | 164700884 | *SI* |  | chr12 | 49448971 | 49449191 | *KMT2D* |
| chr3 | 164704851 | 164705039 | *SI* |  | chr12 | 56992566 | 56992789 | *BAZ2A* |
| chr3 | 164709094 | 164709312 | *SI* |  | chr12 | 56992566 | 56992789 | *BAZ2A* |
| chr3 | 164709914 | 164710329 | *SI* |  | chr12 | 57020797 | 57021025 | *BAZ2A* |
| chr3 | 164711868 | 164712294 | *SI* |  | chr12 | 57020797 | 57021025 | *BAZ2A* |
| chr3 | 164714143 | 164714643 | *SI* |  | chr12 | 69202028 | 69202257 | *MDM2* |
| chr3 | 164716199 | 164716577 | *SI* |  | chr12 | 69202028 | 69202257 | *MDM2* |
| chr3 | 164724490 | 164724771 | *SI* |  | chr12 | 69202554 | 69202784 | *MDM2* |
| chr3 | 164725519 | 164725839 | *SI* |  | chr12 | 69202554 | 69202784 | *MDM2* |
| chr3 | 164727005 | 164727199 | *SI* |  | chr12 | 69205139 | 69205356 | *MDM2* |
| chr3 | 164730665 | 164730868 | *SI* |  | chr12 | 69205139 | 69205356 | *MDM2* |
| chr3 | 164732838 | 164733060 | *SI* |  | chr12 | 69207783 | 69207931 | *MDM2* |
| chr3 | 164733581 | 164733945 | *SI* |  | chr12 | 69207783 | 69207931 | *MDM2* |
| chr3 | 164735180 | 164735936 | *SI* |  | chr12 | 69211977 | 69212182 | *MDM2* |
| chr3 | 164737255 | 164737670 | *SI* |  | chr12 | 69211977 | 69212182 | *MDM2* |
| chr3 | 164738848 | 164739192 | *SI* |  | chr12 | 69214492 | 69214713 | *MDM2* |
| chr3 | 164741205 | 164741569 | *SI* |  | chr12 | 69214492 | 69214713 | *MDM2* |
| chr3 | 164748468 | 164748751 | *SI* |  | chr12 | 69217011 | 69217235 | *MDM2* |
| chr3 | 164750219 | 164750560 | *SI* |  | chr12 | 69217011 | 69217235 | *MDM2* |
| chr3 | 164751107 | 164751277 | *SI* |  | chr12 | 69217750 | 69217970 | *MDM2* |
| chr3 | 164754135 | 164754319 | *SI* |  | chr12 | 69217750 | 69217970 | *MDM2* |
| chr3 | 164755598 | 164755896 | *SI* |  | chr12 | 69218015 | 69218186 | *MDM2* |
| chr3 | 164756682 | 164756984 | *SI* |  | chr12 | 69218015 | 69218186 | *MDM2* |
| chr3 | 164757590 | 164757809 | *SI* |  | chr12 | 69219467 | 69219684 | *MDM2* |
| chr3 | 164758616 | 164759083 | *SI* |  | chr12 | 69219467 | 69219684 | *MDM2* |
| chr3 | 164760838 | 164760991 | *SI* |  | chr12 | 69222215 | 69222391 | *MDM2* |
| chr3 | 164764457 | 164764873 | *SI* |  | chr12 | 69222215 | 69222391 | *MDM2* |
| chr3 | 164766721 | 164767151 | *SI* |  | chr12 | 69222438 | 69222656 | *MDM2* |
| chr3 | 164767443 | 164767710 | *SI* |  | chr12 | 69222438 | 69222656 | *MDM2* |
| chr3 | 164772844 | 164773250 | *SI* |  | chr12 | 69224229 | 69224442 | *MDM2* |
| chr3 | 164776653 | 164777118 | *SI* |  | chr12 | 69224229 | 69224442 | *MDM2* |
| chr3 | 164777535 | 164777861 | *SI* |  | chr12 | 69225673 | 69225901 | *MDM2* |
| chr3 | 164780027 | 164780431 | *SI* |  | chr12 | 69225673 | 69225901 | *MDM2* |
| chr3 | 164781028 | 164781432 | *SI* |  | chr12 | 69230784 | 69231017 | *MDM2* |
| chr3 | 164782870 | 164783290 | *SI* |  | chr12 | 69230784 | 69231017 | *MDM2* |
| chr3 | 164784976 | 164785284 | *SI* |  | chr12 | 69231497 | 69231677 | *MDM2* |
| chr3 | 164786477 | 164786635 | *SI* |  | chr12 | 69231497 | 69231677 | *MDM2* |
| chr3 | 164786824 | 164787025 | *SI* |  | chr12 | 69234934 | 69235134 | *MDM2* |
| chr3 | 164792159 | 164792581 | *SI* |  | chr12 | 69234934 | 69235134 | *MDM2* |
| chr3 | 164793641 | 164793892 | *SI* |  | chr13 | 50606926 | 50607099 | *DLEU2* |
| chr4 | 126237442 | 126242845 | *FAT4* |  | chr13 | 50606926 | 50607099 | *DLEU2* |
| chr4 | 126319887 | 126320096 | *FAT4* |  | chr13 | 50610090 | 50610316 | *DLEU2* |
| chr4 | 126327876 | 126328301 | *FAT4* |  | chr13 | 50610090 | 50610316 | *DLEU2* |
| chr4 | 126329469 | 126330015 | *FAT4* |  | chr13 | 50610382 | 50610612 | *DLEU2* |
| chr4 | 126335999 | 126337024 | *FAT4* |  | chr13 | 50610382 | 50610612 | *DLEU2* |
| chr4 | 126337582 | 126337798 | *FAT4* |  | chr13 | 50613507 | 50613714 | *DLEU2* |
| chr4 | 126355217 | 126355679 | *FAT4* |  | chr13 | 50613507 | 50613714 | *DLEU2* |
| chr4 | 126367280 | 126367703 | *FAT4* |  | chr13 | 50614578 | 50614800 | *DLEU2* |
| chr4 | 126369600 | 126373970 | *FAT4* |  | chr13 | 50614578 | 50614800 | *DLEU2* |
| chr4 | 126384651 | 126384861 | *FAT4* |  | chr13 | 50617451 | 50617672 | *DLEU2* |
| chr4 | 126389659 | 126390108 | *FAT4* |  | chr13 | 50617451 | 50617672 | *DLEU2* |
| chr4 | 126397168 | 126397565 | *FAT4* |  | chr13 | 50621253 | 50621460 | *DLEU2* |
| chr4 | 126398164 | 126398533 | *FAT4* |  | chr13 | 50621253 | 50621460 | *DLEU2* |
| chr4 | 126400723 | 126401138 | *FAT4* |  | chr13 | 50623991 | 50624207 | *DLEU2* |
| chr4 | 126402645 | 126402962 | *FAT4* |  | chr13 | 50623991 | 50624207 | *DLEU2* |
| chr4 | 126408404 | 126408766 | *FAT4* |  | chr13 | 50626569 | 50626786 | *DLEU2* |
| chr4 | 126410917 | 126412964 | *FAT4* |  | chr13 | 50626569 | 50626786 | *DLEU2* |
| chr4 | 153243999 | 153244419 | *FBXW7* |  | chr13 | 50630689 | 50630908 | *DLEU2* |
| chr4 | 153245192 | 153245597 | *FBXW7* |  | chr13 | 50630689 | 50630908 | *DLEU2* |
| chr4 | 153247080 | 153247499 | *FBXW7* |  | chr13 | 50632828 | 50633050 | *DLEU2* |
| chr4 | 153249344 | 153249559 | *FBXW7* |  | chr13 | 50632828 | 50633050 | *DLEU2* |
| chr4 | 153250777 | 153250988 | *FBXW7* |  | chr13 | 50637576 | 50637799 | *DLEU2* |
| chr4 | 153251814 | 153252032 | *FBXW7* |  | chr13 | 50637576 | 50637799 | *DLEU2* |
| chr4 | 153253617 | 153253976 | *FBXW7* |  | chr13 | 50638400 | 50638623 | *DLEU2* |
| chr4 | 153258906 | 153259125 | *FBXW7* |  | chr13 | 50638400 | 50638623 | *DLEU2* |
| chr4 | 153267930 | 153268306 | *FBXW7* |  | chr13 | 50644272 | 50644492 | *DLEU2* |
| chr4 | 153271101 | 153271316 | *FBXW7* |  | chr13 | 50644272 | 50644492 | *DLEU2* |
| chr4 | 153273526 | 153273888 | *FBXW7* |  | chr13 | 50645035 | 50645231 | *DLEU2* |
| chr4 | 153303306 | 153303522 | *FBXW7* |  | chr13 | 50645035 | 50645231 | *DLEU2* |
| chr4 | 153332321 | 153332960 | *FBXW7* |  | chr13 | 50646102 | 50646320 | *DLEU2* |
| chr4 | 187509589 | 187510379 | *FAT1* |  | chr13 | 50646102 | 50646320 | *DLEU2* |
| chr4 | 187516782 | 187517008 | *FAT1* |  | chr13 | 50648383 | 50648564 | *DLEU2* |
| chr4 | 187517680 | 187518330 | *FAT1* |  | chr13 | 50648383 | 50648564 | *DLEU2* |
| chr4 | 187518778 | 187519002 | *FAT1* |  | chr13 | 50655843 | 50656069 | *DLEU2* |
| chr4 | 187519033 | 187519312 | *FAT1* |  | chr13 | 50655843 | 50656069 | *DLEU2* |
| chr4 | 187520944 | 187521519 | *FAT1* |  | chr13 | 104177412 | 104177632 | *-* |
| chr4 | 187522403 | 187522723 | *FAT1* |  | chr13 | 104177412 | 104177632 | *-* |
| chr4 | 187524017 | 187524231 | *FAT1* |  | chr13 | 104182574 | 104182792 | *-* |
| chr4 | 187524311 | 187525144 | *FAT1* |  | chr13 | 104182574 | 104182792 | *-* |
| chr4 | 187525320 | 187525744 | *FAT1* |  | chr13 | 104187557 | 104187780 | *-* |
| chr4 | 187527184 | 187527397 | *FAT1* |  | chr13 | 104187557 | 104187780 | *-* |
| chr4 | 187530283 | 187530506 | *FAT1* |  | chr13 | 104189340 | 104189567 | *-* |
| chr4 | 187530781 | 187531174 | *FAT1* |  | chr13 | 104189340 | 104189567 | *-* |
| chr4 | 187532509 | 187532918 | *FAT1* |  | chr13 | 104190208 | 104190429 | *-* |
| chr4 | 187534222 | 187534507 | *FAT1* |  | chr13 | 104190208 | 104190429 | *-* |
| chr4 | 187535217 | 187535514 | *FAT1* |  | chr13 | 104191479 | 104191696 | *-* |
| chr4 | 187538079 | 187538477 | *FAT1* |  | chr13 | 104191479 | 104191696 | *-* |
| chr4 | 187538721 | 187543051 | *FAT1* |  | chr13 | 104195226 | 104195441 | *-* |
| chr4 | 187549096 | 187549523 | *FAT1* |  | chr13 | 104195226 | 104195441 | *-* |
| chr4 | 187549554 | 187549973 | *FAT1* |  | chr13 | 104197100 | 104197321 | *-* |
| chr4 | 187554801 | 187555014 | *FAT1* |  | chr13 | 104197100 | 104197321 | *-* |
| chr4 | 187557071 | 187557408 | *FAT1* |  | chr13 | 104197369 | 104197594 | *-* |
| chr4 | 187557577 | 187558111 | *FAT1* |  | chr13 | 104197369 | 104197594 | *-* |
| chr4 | 187560811 | 187560998 | *FAT1* |  | chr13 | 104198617 | 104198844 | *-* |
| chr4 | 187584427 | 187584776 | *FAT1* |  | chr13 | 104198617 | 104198844 | *-* |
| chr4 | 187627638 | 187630994 | *FAT1* |  | chr15 | 93444386 | 93444609 | *CHD2* |
| chr5 | 15500755 | 15500928 | *FBXL7* |  | chr15 | 93467516 | 93467647 | *CHD2* |
| chr5 | 15616017 | 15616239 | *FBXL7* |  | chr15 | 93467516 | 93467787 | *CHD2* |
| chr5 | 15927879 | 15928327 | *FBXL7* |  | chr15 | 93470383 | 93470598 | *CHD2* |
| chr5 | 15928454 | 15928618 | *FBXL7* |  | chr15 | 93470383 | 93470598 | *CHD2* |
| chr5 | 15936415 | 15937314 | *FBXL7* |  | chr15 | 93472196 | 93472388 | *CHD2* |
| chr5 | 150884936 | 150885418 | *FAT2* |  | chr15 | 93480679 | 93480898 | *CHD2* |
| chr5 | 150885440 | 150885666 | *FAT2* |  | chr15 | 93480679 | 93480898 | *CHD2* |
| chr5 | 150886646 | 150887201 | *FAT2* |  | chr15 | 93482786 | 93482970 | *CHD2* |
| chr5 | 150889544 | 150889773 | *FAT2* |  | chr15 | 93484865 | 93485082 | *CHD2* |
| chr5 | 150891692 | 150892173 | *FAT2* |  | chr15 | 93484865 | 93485288 | *CHD2* |
| chr5 | 150897159 | 150897370 | *FAT2* |  | chr15 | 93485927 | 93486102 | *CHD2* |
| chr5 | 150900829 | 150901652 | *FAT2* |  | chr15 | 93485927 | 93486309 | *CHD2* |
| chr5 | 150905233 | 150905534 | *FAT2* |  | chr15 | 93487580 | 93487793 | *CHD2* |
| chr5 | 150906750 | 150906967 | *FAT2* |  | chr15 | 93487580 | 93487793 | *CHD2* |
| chr5 | 150907529 | 150907757 | *FAT2* |  | chr15 | 93488928 | 93489487 | *CHD2* |
| chr5 | 150908704 | 150909055 | *FAT2* |  | chr15 | 93489111 | 93489330 | *CHD2* |
| chr5 | 150911122 | 150911642 | *FAT2* |  | chr15 | 93492123 | 93492340 | *CHD2* |
| chr5 | 150913894 | 150914209 | *FAT2* |  | chr15 | 93492123 | 93492340 | *CHD2* |
| chr5 | 150917323 | 150917546 | *FAT2* |  | chr15 | 93496401 | 93496808 | *CHD2* |
| chr5 | 150920095 | 150920340 | *FAT2* |  | chr15 | 93496611 | 93496808 | *CHD2* |
| chr5 | 150921673 | 150925903 | *FAT2* |  | chr15 | 93498471 | 93498754 | *CHD2* |
| chr5 | 150928809 | 150929096 | *FAT2* |  | chr15 | 93498682 | 93498754 | *CHD2* |
| chr5 | 150930089 | 150930490 | *FAT2* |  | chr15 | 93499682 | 93499889 | *CHD2* |
| chr5 | 150930991 | 150931204 | *FAT2* |  | chr15 | 93510528 | 93510751 | *CHD2* |
| chr5 | 150932658 | 150932960 | *FAT2* |  | chr15 | 93510528 | 93510751 | *CHD2* |
| chr5 | 150933823 | 150934241 | *FAT2* |  | chr15 | 93514828 | 93515188 | *CHD2* |
| chr5 | 150935806 | 150936022 | *FAT2* |  | chr15 | 93515033 | 93515188 | *CHD2* |
| chr5 | 150942848 | 150943205 | *FAT2* |  | chr15 | 93515359 | 93515676 | *CHD2* |
| chr5 | 150945193 | 150947930 | *FAT2* |  | chr15 | 93515520 | 93515676 | *CHD2* |
| chr5 | 150948121 | 150948507 | *FAT2* |  | chr15 | 93518026 | 93518241 | *CHD2* |
| chr6 | 393004 | 393526 | *IRF4* |  | chr15 | 93521439 | 93521664 | *CHD2* |
| chr6 | 394802 | 395023 | *IRF4* |  | chr15 | 93521439 | 93521664 | *CHD2* |
| chr6 | 395806 | 396016 | *IRF4* |  | chr15 | 93522300 | 93522526 | *CHD2* |
| chr6 | 397110 | 397292 | *IRF4* |  | chr15 | 93523842 | 93524062 | *CHD2* |
| chr6 | 398801 | 399006 | *IRF4* |  | chr15 | 93523842 | 93524203 | *CHD2* |
| chr6 | 401247 | 401784 | *IRF4* |  | chr15 | 93524548 | 93524736 | *CHD2* |
| chr6 | 404964 | 405183 | *IRF4* |  | chr15 | 93524548 | 93524736 | *CHD2* |
| chr6 | 407395 | 407604 | *IRF4* |  | chr15 | 93527547 | 93527761 | *CHD2* |
| chr6 | 26156451 | 26157120 | *HIST1H1E* |  | chr15 | 93528701 | 93528928 | *CHD2* |
| chr6 | 26157205 | 26157285 | *HIST1H1E* |  | chr15 | 93528701 | 93528928 | *CHD2* |
| chr6 | 44226873 | 44227061 | *NFKBIE* |  | chr15 | 93534559 | 93534753 | *CHD2* |
| chr6 | 44227615 | 44228024 | *NFKBIE* |  | chr15 | 93536043 | 93536267 | *CHD2* |
| chr6 | 44228093 | 44228308 | *NFKBIE* |  | chr15 | 93536043 | 93536267 | *CHD2* |
| chr6 | 44229163 | 44229709 | *NFKBIE* |  | chr15 | 93540133 | 93540352 | *CHD2* |
| chr6 | 44230269 | 44230458 | *NFKBIE* |  | chr15 | 93540444 | 93540639 | *CHD2* |
| chr6 | 44232636 | 44233244 | *NFKBIE* |  | chr15 | 93540444 | 93540639 | *CHD2* |
| chr6 | 44233261 | 44233547 | *NFKBIE* |  | chr15 | 93541680 | 93541900 | *CHD2* |
| chr7 | 82387687 | 82388104 | *PCLO* |  | chr15 | 93543693 | 93543912 | *CHD2* |
| chr7 | 82389765 | 82390108 | *PCLO* |  | chr15 | 93543693 | 93543912 | *CHD2* |
| chr7 | 82390591 | 82390905 | *PCLO* |  | chr15 | 93545347 | 93545574 | *CHD2* |
| chr7 | 82430739 | 82430951 | *PCLO* |  | chr15 | 93547806 | 93548022 | *CHD2* |
| chr7 | 82434963 | 82435184 | *PCLO* |  | chr15 | 93547806 | 93548022 | *CHD2* |
| chr7 | 82451726 | 82452044 | *PCLO* |  | chr15 | 93552362 | 93552576 | *CHD2* |
| chr7 | 82453510 | 82453844 | *PCLO* |  | chr15 | 93555379 | 93555587 | *CHD2* |
| chr7 | 82455736 | 82456152 | *PCLO* |  | chr15 | 93555379 | 93555756 | *CHD2* |
| chr7 | 82457124 | 82457338 | *PCLO* |  | chr15 | 93557730 | 93557955 | *CHD2* |
| chr7 | 82464909 | 82465115 | *PCLO* |  | chr15 | 93557730 | 93558144 | *CHD2* |
| chr7 | 82467349 | 82467780 | *PCLO* |  | chr15 | 93563211 | 93563429 | *CHD2* |
| chr7 | 82470718 | 82470882 | *PCLO* |  | chr15 | 93563211 | 93563523 | *CHD2* |
| chr7 | 82474551 | 82474831 | *PCLO* |  | chr15 | 93567503 | 93567662 | *CHD2* |
| chr7 | 82475833 | 82476050 | *PCLO* |  | chr15 | 93567503 | 93567940 | *CHD2* |
| chr7 | 82476391 | 82476612 | *PCLO* |  | chr15 | 93567853 | 93567940 | *CHD2* |
| chr7 | 82508553 | 82508816 | *PCLO* |  | chr16 | 81946082 | 81946303 | *PLCG2* |
| chr7 | 82531980 | 82532081 | *PCLO* |  | chr16 | 81946082 | 81946303 | *PLCG2* |
| chr7 | 82538151 | 82538371 | *PCLO* |  | chr16 | 81952960 | 81953180 | *PLCG2* |
| chr7 | 82543917 | 82546283 | *PCLO* |  | chr16 | 81952960 | 81953180 | *PLCG2* |
| chr7 | 82578787 | 82580811 | *PCLO* |  | chr16 | 81962016 | 81962236 | *PLCG2* |
| chr7 | 82581127 | 82586279 | *PCLO* |  | chr16 | 81962016 | 81962236 | *PLCG2* |
| chr7 | 82595069 | 82595886 | *PCLO* |  | chr17 | 7572853 | 7573081 | *TP53* |
| chr7 | 82763497 | 82764981 | *PCLO* |  | chr17 | 7573878 | 7574094 | *TP53* |
| chr7 | 82783885 | 82784537 | *PCLO* |  | chr17 | 7573878 | 7574094 | *TP53* |
| chr7 | 82784561 | 82785823 | *PCLO* |  | chr17 | 7576495 | 7576760 | *TP53* |
| chr7 | 82791654 | 82791885 | *PCLO* |  | chr17 | 7576559 | 7576760 | *TP53* |
| chr7 | 82791903 | 82792072 | *PCLO* |  | chr17 | 7576783 | 7577196 | *TP53* |
| chr7 | 124463961 | 124464182 | *POT1* |  | chr17 | 7576972 | 7577196 | *TP53* |
| chr7 | 124465155 | 124465510 | *POT1* |  | chr17 | 7577398 | 7577615 | *TP53* |
| chr7 | 124467185 | 124467375 | *POT1* |  | chr17 | 7578087 | 7578296 | *TP53* |
| chr7 | 124469129 | 124469521 | *POT1* |  | chr17 | 7578087 | 7578560 | *TP53* |
| chr7 | 124475276 | 124475499 | *POT1* |  | chr17 | 7578385 | 7578560 | *TP53* |
| chr7 | 124481014 | 124481250 | *POT1* |  | chr17 | 7579288 | 7579519 | *TP53* |
| chr7 | 124482675 | 124483051 | *POT1* |  | chr17 | 7579404 | 7579519 | *TP53* |
| chr7 | 124486939 | 124487113 | *POT1* |  | chr17 | 7579610 | 7579966 | *TP53* |
| chr7 | 124491738 | 124492054 | *POT1* |  | chr17 | 7579738 | 7579966 | *TP53* |
| chr7 | 124492923 | 124493311 | *POT1* |  | chr17 | 37947584 | 37947806 | *IKZF3* |
| chr7 | 124498830 | 124499176 | *POT1* |  | chr17 | 37947584 | 37947806 | *IKZF3* |
| chr7 | 124503331 | 124503725 | *POT1* |  | chr19 | 1440335 | 1440565 | *RPS15* |
| chr7 | 124510779 | 124511153 | *POT1* |  | chr19 | 1440335 | 1440565 | *RPS15* |
| chr7 | 124532293 | 124532506 | *POT1* |  | chr19 | 50922176 | 50922391 | *SPIB* |
| chr7 | 124537139 | 124537305 | *POT1* |  | chr19 | 50923158 | 50923281 | *SPIB* |
| chr7 | 140434378 | 140434640 | *BRAF* |  | chr19 | 50923158 | 50923281 | *SPIB* |
| chr7 | 140439596 | 140439801 | *BRAF* |  | chr19 | 50925649 | 50925876 | *SPIB* |
| chr7 | 140449055 | 140449273 | *BRAF* |  | chr19 | 50926061 | 50926276 | *SPIB* |
| chr7 | 140453024 | 140453240 | *BRAF* |  | chr19 | 50926061 | 50926306 | *SPIB* |
| chr7 | 140453842 | 140454071 | *BRAF* |  | chr19 | 50926811 | 50926922 | *SPIB* |
| chr7 | 140476597 | 140476901 | *BRAF* |  | chr19 | 50926811 | 50927076 | *SPIB* |
| chr7 | 140477603 | 140477880 | *BRAF* |  | chr19 | 50931202 | 50931413 | *SPIB* |
| chr7 | 140481301 | 140481520 | *BRAF* |  | chr19 | 50931202 | 50932794 | *SPIB* |
| chr7 | 140482785 | 140482988 | *BRAF* |  | chr19 | 50931537 | 50931684 | *SPIB* |
| chr7 | 140487253 | 140487438 | *BRAF* |  | chr19 | 50931815 | 50931965 | *SPIB* |
| chr7 | 140493956 | 140494419 | *BRAF* |  | chr19 | 50932115 | 50932340 | *SPIB* |
| chr7 | 140500104 | 140500309 | *BRAF* |  | chr19 | 50932528 | 50932655 | *SPIB* |
| chr7 | 140501037 | 140501391 | *BRAF* |  | chr19 | 50932813 | 50932958 | *SPIB* |
| chr7 | 140507659 | 140507877 | *BRAF* |  | chr19 | 50932813 | 50932958 | *SPIB* |
| chr7 | 140508608 | 140508819 | *BRAF* |  | chr19 | 50933273 | 50933707 | *SPIB* |
| chr7 | 140534303 | 140534680 | *BRAF* |  | chr19 | 50933482 | 50933707 | *SPIB* |
| chr7 | 140549800 | 140550073 | *BRAF* |  | chr19 | 50934041 | 50934319 | *SPIB* |
| chr7 | 140624312 | 140624499 | *BRAF* |  | chr19 | 50934183 | 50934319 | *SPIB* |
| chr8 | 113236966 | 113237192 | *CSMD3* |  | chr20 | 35521286 | 35521505 | *SAMHD1* |
| chr8 | 113240875 | 113241246 | *CSMD3* |  | chr20 | 35526172 | 35526390 | *SAMHD1* |
| chr8 | 113243673 | 113243894 | *CSMD3* |  | chr20 | 35526172 | 35526390 | *SAMHD1* |
| chr8 | 113246422 | 113246810 | *CSMD3* |  | chr20 | 35526754 | 35526977 | *SAMHD1* |
| chr8 | 113249273 | 113249671 | *CSMD3* |  | chr20 | 35532513 | 35532699 | *SAMHD1* |
| chr8 | 113253903 | 113254098 | *CSMD3* |  | chr20 | 35532513 | 35532699 | *SAMHD1* |
| chr8 | 113256552 | 113256977 | *CSMD3* |  | chr20 | 35533757 | 35533975 | *SAMHD1* |
| chr8 | 113259074 | 113259394 | *CSMD3* |  | chr20 | 35539600 | 35539813 | *SAMHD1* |
| chr8 | 113266466 | 113266680 | *CSMD3* |  | chr20 | 35539600 | 35539813 | *SAMHD1* |
| chr8 | 113267305 | 113267710 | *CSMD3* |  | chr20 | 35540816 | 35541009 | *SAMHD1* |
| chr8 | 113275791 | 113276049 | *CSMD3* |  | chr20 | 35545097 | 35545314 | *SAMHD1* |
| chr8 | 113277630 | 113277829 | *CSMD3* |  | chr20 | 35545097 | 35545473 | *SAMHD1* |
| chr8 | 113293221 | 113293610 | *CSMD3* |  | chr20 | 35547731 | 35547953 | *SAMHD1* |
| chr8 | 113299165 | 113299480 | *CSMD3* |  | chr20 | 35547731 | 35547953 | *SAMHD1* |
| chr8 | 113301588 | 113301801 | *CSMD3* |  | chr20 | 35555477 | 35555670 | *SAMHD1* |
| chr8 | 113303705 | 113304057 | *CSMD3* |  | chr20 | 35558951 | 35559170 | *SAMHD1* |
| chr8 | 113304702 | 113305108 | *CSMD3* |  | chr20 | 35558951 | 35559283 | *SAMHD1* |
| chr8 | 113307926 | 113308290 | *CSMD3* |  | chr20 | 35563374 | 35563450 | *SAMHD1* |
| chr8 | 113313826 | 113314218 | *CSMD3* |  | chr20 | 35563374 | 35563625 | *SAMHD1* |
| chr8 | 113316753 | 113317149 | *CSMD3* |  | chr20 | 35569237 | 35569457 | *SAMHD1* |
| chr8 | 113318223 | 113318433 | *CSMD3* |  | chr20 | 35569237 | 35569657 | *SAMHD1* |
| chr8 | 113323065 | 113323436 | *CSMD3* |  | chr20 | 35575078 | 35575270 | *SAMHD1* |
| chr8 | 113326025 | 113326365 | *CSMD3* |  | chr20 | 35575078 | 35575270 | *SAMHD1* |
| chr8 | 113326579 | 113326860 | *CSMD3* |  | chr20 | 35579850 | 35580066 | *SAMHD1* |
| chr8 | 113330864 | 113331196 | *CSMD3* |  | chrX | 41193469 | 41193635 | *DDX3X* |
| chr8 | 113332037 | 113332241 | *CSMD3* |  | chrX | 41196539 | 41196753 | *DDX3X* |
| chr8 | 113347527 | 113347736 | *CSMD3* |  | chrX | 41198205 | 41198421 | *DDX3X* |
| chr8 | 113348760 | 113349095 | *CSMD3* |  | chrX | 41200558 | 41200907 | *DDX3X* |
| chr8 | 113349590 | 113349985 | *CSMD3* |  | chrX | 41201594 | 41201912 | *DDX3X* |
| chr8 | 113353526 | 113353929 | *CSMD3* |  | chrX | 41201945 | 41202163 | *DDX3X* |
| chr8 | 113358321 | 113358451 | *CSMD3* |  | chrX | 41202301 | 41202673 | *DDX3X* |
| chr8 | 113363266 | 113363479 | *CSMD3* |  | chrX | 41202947 | 41203123 | *DDX3X* |
| chr8 | 113364483 | 113364866 | *CSMD3* |  | chrX | 41203143 | 41203686 | *DDX3X* |
| chr8 | 113392564 | 113392702 | *CSMD3* |  | chrX | 41204260 | 41204869 | *DDX3X* |
| chr8 | 113395734 | 113395938 | *CSMD3* |  | chrX | 41205355 | 41205701 | *DDX3X* |
| chr8 | 113402814 | 113403032 | *CSMD3* |  | chrX | 41205741 | 41205913 | *DDX3X* |
| chr8 | 113418593 | 113418991 | *CSMD3* |  | chrX | 41206084 | 41206293 | *DDX3X* |
| chr8 | 113420345 | 113420770 | *CSMD3* |  | chrX | 41206544 | 41206721 | *DDX3X* |
| chr8 | 113421125 | 113421284 | *CSMD3* |  | chrX | 41206819 | 41207042 | *DDX3X* |
| chr8 | 113484786 | 113485125 | *CSMD3* |  | chrX | 70338452 | 70338799 | *MED12* |
| chr8 | 113504528 | 113504936 | *CSMD3* |  | chrX | 70339147 | 70339367 | *MED12* |
| chr8 | 113515816 | 113516216 | *CSMD3* |  | chrX | 70339527 | 70339735 | *MED12* |
| chr8 | 113518722 | 113519124 | *CSMD3* |  | chrX | 70339825 | 70340052 | *MED12* |
| chr8 | 113529138 | 113529605 | *CSMD3* |  | chrX | 70340810 | 70341029 | *MED12* |
| chr8 | 113562776 | 113563163 | *CSMD3* |  | chrX | 70341122 | 70341684 | *MED12* |
| chr8 | 113564788 | 113564973 | *CSMD3* |  | chrX | 70342002 | 70342225 | *MED12* |
| chr8 | 113568799 | 113569200 | *CSMD3* |  | chrX | 70342320 | 70342506 | *MED12* |
| chr8 | 113585715 | 113585916 | *CSMD3* |  | chrX | 70342545 | 70342765 | *MED12* |
| chr8 | 113599183 | 113599518 | *CSMD3* |  | chrX | 70342896 | 70343119 | *MED12* |
| chr8 | 113648893 | 113649268 | *CSMD3* |  | chrX | 70343400 | 70343622 | *MED12* |
| chr8 | 113650784 | 113651168 | *CSMD3* |  | chrX | 70343839 | 70344246 | *MED12* |
| chr8 | 113657273 | 113657492 | *CSMD3* |  | chrX | 70344535 | 70344740 | *MED12* |
| chr8 | 113662219 | 113662585 | *CSMD3* |  | chrX | 70344798 | 70345024 | *MED12* |
| chr8 | 113668356 | 113668713 | *CSMD3* |  | chrX | 70345152 | 70345382 | *MED12* |
| chr8 | 113678355 | 113678709 | *CSMD3* |  | chrX | 70345448 | 70345629 | *MED12* |
| chr8 | 113694603 | 113694886 | *CSMD3* |  | chrX | 70345855 | 70346056 | *MED12* |
| chr8 | 113697498 | 113698089 | *CSMD3* |  | chrX | 70346139 | 70346367 | *MED12* |
| chr8 | 113702069 | 113702297 | *CSMD3* |  | chrX | 70346780 | 70347005 | *MED12* |
| chr8 | 113812312 | 113812528 | *CSMD3* |  | chrX | 70347128 | 70347355 | *MED12* |
| chr8 | 113841826 | 113842050 | *CSMD3* |  | chrX | 70347583 | 70348008 | *MED12* |
| chr8 | 113871316 | 113871639 | *CSMD3* |  | chrX | 70348138 | 70348365 | *MED12* |
| chr8 | 113933694 | 113934012 | *CSMD3* |  | chrX | 70348383 | 70348611 | *MED12* |
| chr8 | 113959922 | 113960215 | *CSMD3* |  | chrX | 70348865 | 70349089 | *MED12* |
| chr8 | 113966859 | 113967053 | *CSMD3* |  | chrX | 70349110 | 70349335 | *MED12* |
| chr8 | 113988047 | 113988384 | *CSMD3* |  | chrX | 70349510 | 70350113 | *MED12* |
| chr8 | 114031249 | 114031455 | *CSMD3* |  | chrX | 70351322 | 70351548 | *MED12* |
| chr8 | 114110948 | 114111173 | *CSMD3* |  | chrX | 70351882 | 70352106 | *MED12* |
| chr8 | 114185940 | 114186154 | *CSMD3* |  | chrX | 70352121 | 70352448 | *MED12* |
| chr8 | 114290813 | 114291035 | *CSMD3* |  | chrX | 70352491 | 70352863 | *MED12* |
| chr8 | 114326736 | 114327089 | *CSMD3* |  | chrX | 70352937 | 70353162 | *MED12* |
| chr8 | 114388863 | 114389082 | *CSMD3* |  | chrX | 70354167 | 70354388 | *MED12* |
| chr8 | 114448743 | 114449164 | *CSMD3* |  | chrX | 70354516 | 70354730 | *MED12* |
| chr9 | 21968186 | 21968336 | *CDKN2A* |  | chrX | 70354919 | 70355144 | *MED12* |
| chr9 | 21968614 | 21968825 | *CDKN2A* |  | chrX | 70355986 | 70356520 | *MED12* |
| chr9 | 21970770 | 21970975 | *CDKN2A* |  | chrX | 70356704 | 70356893 | *MED12* |
| chr9 | 21970985 | 21971212 | *CDKN2A* |  | chrX | 70356975 | 70357301 | *MED12* |
| chr9 | 21974466 | 21974699 | *CDKN2A* |  | chrX | 70357368 | 70357798 | *MED12* |
| chr9 | 21994065 | 21994411 | *CDKN2A* |  | chrX | 70360291 | 70360714 | *MED12* |
| chr9 | 37034057 | 37034282 | *PAX5* |  | chrX | 70361075 | 70361295 | *MED12* |
| chr9 | 37369346 | 37369568 | *PAX5* |  | chrX | 70361682 | 70361874 | *MED12* |
| chr9 | 37369610 | 37369817 | *PAX5* |  | chrX | 70361894 | 70362120 | *MED12* |
| chr9 | 37371096 | 37371327 | *PAX5* |  | chrX | 100611029 | 100611256 | *BTK* |
