## Supplementary Table 2 for "*SAMHD1* mutations in mantle cell lymphoma are recurrent and confer *in vitro* resistance to nucleoside analogues"

List of all found mutations with the custom next generation sequencing panels. CDS indicates coding sequence changes, AA indicates amino acid change and VAF indicates variant allele frequency.

| **Patient ID** | **Cohort** | **Gene** | **CDS** | **AA** | **VAF (%)** |
| --- | --- | --- | --- | --- | --- |
| Pat 1 | Blood | *FAT1* | c.4291G>A | p.A1431T | 53.35 |
|  |  | *TP53* | c.421_422insTGC | p.R141delinsLR | 12.96 |
| Pat 2 | Blood | *RYR2* | c.8003G>A | p.C2668Y | 21.78 |
| Pat 3 | Blood | *TP53* | c.451C>A | p.P151T | 90.48 |
| Pat 4 | Blood | *FAT1* | c.9923C>A | p.T3308N | 11.85 |
| Pat 5 | Blood | *ATM* | c.67C>T | p.R23X | 53.01 |
|  |  | *ATM* | c.4324T>C | p.Y1442H | 44.34 |
|  |  | *SAMHD1* | c.997C>T | p.R333C | 99 |
|  |  | *TP53* | c.421C>T | p.R141C | 53.85 |
| Pat 6 | Blood | *NOTCH1* | 3'UTR | - | 31.11 |
|  |  | *TP53* | c.438delT | p.P146fs | 22.99 |
| Pat 7 | Blood | *BIRC3* | c.1663A>T | p.R555X | 12.87 |
| Pat 8 | Blood | *BIRC3* | c.1669T>G | p.C557G | 86.89 |
| Pat 9 | Blood | *NOTCH1* | c.7330C>T | p.Q2444X | 58.33 |
|  |  | *TP53* | c.422G>A | p.R141H | 82.73 |
| Pat 10 | Blood | *-* | - | - | - |
| Pat 11 | Blood | *TP53* | c.263A>G | p.Y88C | 82.47 |
| Pat 12 | Blood | *ATM* | c.1375_1377del | p.459_459del | 50 |
|  |  | *ATM* | c.6115G>A | p.E2039K | 33.33 |
| Pat 13 | Blood | *-* | - | - | - |
| Pat 14 | Blood | *TP53* | c.783-1G>T | splice site | 94.72 |
| Pat 15 | Blood | *FAT1* | c.11770C>T | p.R3924C | 32.89 |
|  |  | *FAT4* | c.11274delG | p.E3758fs | 39.12 |
| Pat 16 | Blood | *ATM* | c.380delC | p.T127fs | 90.54 |
|  |  | *KMT2D* | c.779dupT | p.L260fs | 48.29 |
|  |  | *NFKBIE* | c.760_763del | p.Y254fs | 56.98 |
| Pat 17 | Blood | *KMT2D* | c.7379delG | p.R2460fs | 44.44 |
| Pat 18 | Blood | *CHD2* | c.3881A>T | p.D1294V | 13.88 |
|  |  | *CHD2* | c.4165dupA | p.M1388fs | 13.66 |
|  |  | *SAMHD1* | c.1181A>G | p.D394G | 96.55 |
| Pat 19 | Blood | *KMT2D* | c.12815dupG | p.G4272fs | 30.45 |
|  |  | *SAMHD1* | c.1456delA | p.K486fs | 94.13 |
| Pat 20 | Blood | *NFKBIE* | c.760_763del | p.Y254fs | 35.57 |
| Pat 21 | Blood | *-* | - | - | - |
| Pat 22 | Blood | *TP53* | c.422G>A | p.R141H | 55.56 |
| Pat 23 | Blood | *ATM* | c.7009T>C | p.C2337R | 60.64 |
|  |  | *HIST1H1E* | c.113C>T | p.P38L | 35.02 |
| Pat 24 | Blood | *TP53* | c.413T>C | p.F138S | 26.89 |
| Pat 25 | Blood | *SAMHD1* | c.22C>T | p.Q8X | 87.29 |
| Pat 26 | Tissue | *TP53* | c.742C>T | p.Arg248Trp | 84.37 |
|  |  | *BRAF* | c.1406G>T | p.Gly469Val | 35.98 |
| Pat 27 | Tissue | *NOTCH1* | c.*371A>G | 3' UTR variant | 80.04 |
| Pat 28 | Tissue | *TP53* | c.767C>A | p.Thr256Lys | 95.97 |
| Pat 29 | Tissue | *ATM* | c.2776A>T | p.Lys926* | 25.36 |
|  |  | *ATM* | c.7397delC | p.Ala2466fs | 31.61 |
|  |  | *SI* | c.5279G>T | p.Gly1760Val | 59.27 |
| Pat 30 | Tissue | *SPEN* | c.2372_2373insTATCA | p.Asp792fs | 33.73 |
|  |  | *NFKBIE* | c.1142_1143insCTGAA | p.Lys381fs | 38.35 |
|  |  | *NFKBIE* | c.506_507dupCC | p.Glu170fs | 33.05 |
|  |  | *ATM* | c.5944C>T | p.Gln1982* | 32.91 |
|  |  | *ATM* | c.9023G>A | p.Arg3008His | 34.03 |
| Pat 31 | Tissue | *ATM* | c.3115delA | p.Arg1039fs | 67.17 |
|  |  | *FAT1* | c.2923T>C | p.Tyr975His | 52.35 |
| Pat 32 | Tissue | *SPEN* | c.10396A>G | p.Ser3466Gly | 27.47 |
|  |  | *ATM* | c.7330G>A | p.Glu2444Lys | 100 |
| Pat 33 | Tissue | *-* | - | - | - |
| Pat 34 | Tissue | *TP53* | c.695T>G | p.Ile232Ser | 26.92 |
|  |  | *SPEN* | c.8069_8070insAA | p.Asn2691fs | 39.9 |
|  |  | *CHD2* | c.4575C>A | p.His1525Gln | 24.49 |
|  |  | *CSMD3* | c.9187A>G | p.Thr3063Ala | 11.06 |
|  |  | *CSMD3* | c.2839G>A | p.Glu947Lys | 17.41 |
| Pat 35 | Tissue | *TP53* | c.743G>A | p.Arg248Gln | 60.6 |
|  |  | *SPEN* | c.3229G>A | p.Gly1077Ser | 48.47 |
|  |  | *ATM* | c.4511C>T | p.Thr1504Ile | 48.84 |
| Pat 36 | Tissue | *TP53* | c.659A>G | p.Tyr220Cys | 95.14 |
|  |  | *KMT2D* | c.2887G>A | p.Ala963Thr | 4.01 |
|  |  | *CHD2* | c.694T>C | p.Tyr232His | 32.03 |
|  |  | *CHD2* | c.695A>T | p.Tyr232Phe | 32.03 |
|  |  | *FAT4* | c.13314_13315delTG | p.Gly4439fs | 16.81 |
|  |  | *NOTCH1* | c.3734T>G | p.Val1245Gly | 23.3 |
|  |  | *RYR2* | c.277G>C | p.Val93Leu | 16.47 |
|  |  | *RYR2* | c.281A>C | p.Asp94Ala | 18.23 |
|  |  | *RYR2* | c.6280G>A | p.Gly2094Ser | 33.33 |
| Pat 37 | Tissue | *CHD2* | c.2210A>G | p.Tyr737Cys | 34.12 |
|  |  | *NRAS* | c.38G>A | p.Gly13Asp | 38.56 |
| Pat 38 | Tissue | *-* | - | - | - |
| Pat 39 | Tissue | *ATM* | c.9023G>A | p.Arg3008His | 58.83 |
|  |  | *FBXW7* | c.349_351delGAG | p.Glu117del | 2.21 |
| Pat 40 | Tissue | *TP53* | c.524G>A | p.Arg175His | 48.7 |
|  |  | *DDX3X* | c.1678_1680delCTT | p.Leu560del | 75.65 |
|  |  | *NFKBIE* | c.137delT | p.Ile46fs | 67.34 |
|  |  | *SPI1* | c.605C>T | p.Ser202Leu | 22 |
|  |  | *PCLO* | c.11470C>T | p.Leu3824Phe | 52.22 |
|  |  | *CSMD3* | c.985A>G | p.Thr329Ala | 7.7 |
|  |  | *FAT4* | c.1243C>G | p.Pro415Ala | 47.77 |
|  |  | *FAT1* | c.12079G>A | p.Val4027Ile | 31.39 |
|  |  | *FAT1* | c.8404G>T | p.Ala2802Ser | 34.71 |
| Pat 41 | Tissue | *TP53* | c.469G>T | p.Val157Phe | 41.78 |
|  |  | *KMT2D* | c.3786C>A | p.Asp1262Glu | 44.79 |
|  |  | *FUBP1* | c.793C>T | p.Arg265Trp | 58.93 |
| Pat 42 | Tissue | *CHD2* | c.5086A>T | p.Arg1696* | 45.44 |
|  |  | *SF3B1* | c.2584G>A | p.Glu862Lys | 58.33 |
| Pat 43 | Tissue | *TP53* | c.857A>T | p.Glu286Val | 79.26 |
|  |  | *SPI1* | c.676C>G | p.Gln226Glu | 34.91 |
|  |  | *NOTCH1* | c.7541_7542delCT | p.Pro2514fs | 78.95 |
| Pat 44 | Tissue | *ATM* | c.5738T>G | p.Val1913Gly | 97.6 |
|  |  | *RYR2* | c.908G>A | p.Gly303Glu | 62.04 |
| Pat 45 | Tissue | *TP53* | c.869G>A | p.Arg290His | 48.75 |
|  |  | *KMT2D* | c.6643T>A | p.Ser2215Thr | 49.41 |
|  |  | *ATM* | c.5599delC | p.Gln1867fs | 61.04 |
|  |  | *ATM* | c.7330G>A | p.Glu2444Lys | 54.26 |
|  |  | *CSMD3* | c.9059A>T | p.Tyr3020Phe | 51.87 |
|  |  | *FAT1* | c.4433T>C | p.Ile1478Thr | 51.25 |
| Pat 46 | Tissue | *SPEN* | c.6994C>T | p.Arg2332Cys | 32.59 |
|  |  | *HIST1H1E* | c.127C>T | p.Leu43Phe | 45.28 |
|  |  | *SI* | c.1780T>C | p.Ser594Pro | 51.88 |
|  |  | *RYR2* | c.5920A>C | p.Asn1974His | 66.46 |
| Pat 47 | Tissue | *TP53* | c.814G>T | p.Val272Leu | 73.14 |
