## Supplementary Table 3 for "*SAMHD1* mutations in mantle cell lymphoma are recurrent and confer *in vitro* resistance to nucleoside analogues"

Cell viability after drug exposure to the four tested drugs (cytarabine, fludarabine, doxorubicine and nutlin-3a) in four different concentrations, 1 denotes 100% cell viability.

| **Drug** | **Conc. (µM)** | **Viability (Normalized to internal DMSO control)** | | | | | | | | | | | | | |
| --- | --- | --- | --- | --- | --- | --- | --- | --- | --- | --- | --- | --- | --- | --- | --- |
|  |  | Pat 3 | Pat 11 | Pat 12 | Pat 13 | Pat 14 | Pat 16 | Pat 17 | Pat 18 | Pat 19 | Pat 20 | Pat 21 | Pat 22 | Pat 23 | Pat 25 |
| Cytarabine | 0.0160 | 1.05 | 1.08 | 1.12 | 1.02 | 1.01 | 1.06 | 1.08 | 1.05 | 1.10 | 1.09 | 1.02 | 1.07 | 1.02 | 1.04 |
| Cytarabine | 0.0800 | 0.86 | 1.02 | 1.01 | 0.99 | 0.96 | 0.98 | 1.08 | 1.04 | 1.10 | 1.04 | 0.93 | 0.94 | 1.00 | 0.97 |
| Cytarabine | 0.4000 | 0.60 | 0.86 | 0.75 | 0.81 | 0.83 | 0.53 | 0.87 | 1.00 | 1.34 | 0.83 | 0.89 | 0.88 | 0.83 | 0.94 |
| Cytarabine | 2.0000 | 0.29 | 0.56 | 0.30 | 0.42 | 0.59 | 0.22 | 0.35 | 0.81 | 0.45 | 0.30 | 0.65 | 0.55 | 0.49 | 0.87 |
| Cytarabine | 10.0000 | 0.18 | 0.40 | 0.13 | 0.21 | 0.33 | 0.15 | 0.14 | 0.47 | 0.60 | 0.14 | 0.29 | 0.42 | 0.26 | 0.66 |
| Doxorubicine | 0.0006 | 1.13 | 0.99 | 1.03 | 0.99 | 1.04 | 0.94 | 0.97 | 1.01 | 1.20 | 1.00 | 1.01 | 1.01 | 0.97 | 1.00 |
| Doxorubicine | 0.0032 | 0.86 | 1.01 | 1.04 | 0.90 | 1.02 | 0.93 | 1.00 | 1.05 | 1.07 | 1.02 | 1.01 | 1.06 | 1.09 | 0.99 |
| Doxorubicine | 0.0160 | 0.94 | 0.99 | 0.95 | 0.95 | 0.98 | 0.95 | 1.03 | 1.04 | 0.67 | 0.98 | 1.01 | 1.05 | 1.02 | 0.97 |
| Doxorubicine | 0.0800 | 0.72 | 0.95 | 0.81 | 1.04 | 0.97 | 0.98 | 0.92 | 1.03 | 0.74 | 0.66 | 0.99 | 1.04 | 0.92 | 0.98 |
| Doxorubicine | 0.4000 | 0.13 | 0.44 | 0.18 | 0.62 | 0.34 | 0.59 | 0.17 | 0.85 | 0.80 | 0.08 | 0.73 | 0.93 | 0.50 | 0.83 |
| Fludarabine | 0.0160 | 0.98 | 1.03 | 1.13 | 1.01 | 1.05 | 1.01 | 1.11 | 1.06 | 0.69 | 1.09 | 0.97 | 1.02 | 0.99 | 0.98 |
| Fludarabine | 0.0800 | 0.80 | 1.04 | 0.99 | 0.90 | 0.96 | 1.03 | 1.09 | 1.02 | 1.06 | 0.98 | 0.95 | 1.02 | 0.92 | 0.97 |
| Fludarabine | 0.4000 | 0.32 | 0.96 | 0.84 | 0.53 | 0.95 | 0.81 | 0.78 | 1.01 | 0.92 | 0.60 | 0.60 | 0.78 | 0.65 | 0.91 |
| Fludarabine | 2.0000 | 0.20 | 0.76 | 0.27 | 0.27 | 0.55 | 0.42 | 0.22 | 0.93 | 0.89 | 0.16 | 0.30 | 0.55 | 0.30 | 0.84 |
| Fludarabine | 10.0000 | 0.11 | 0.47 | 0.15 | 0.20 | 0.21 | 0.15 | 0.06 | 0.76 | 0.63 | 0.07 | 0.21 | 0.34 | 0.22 | 0.66 |
| Nutlin-3a | 0.0160 | 0.85 | 0.98 | 0.93 | 0.89 | 0.96 | 1.01 | 1.08 | 0.96 | 1.27 | 0.98 | 0.99 | 1.03 | 0.90 | 0.99 |
| Nutlin-3a | 0.0800 | 0.92 | 0.97 | 0.98 | 1.04 | 1.02 | 0.97 | 1.13 | 0.99 | 1.07 | 1.00 | 0.96 | 0.95 | 0.90 | 0.95 |
| Nutlin-3a | 0.4000 | 1.03 | 1.03 | 0.94 | 1.04 | 1.00 | 0.93 | 1.07 | 0.97 | 1.14 | 0.90 | 0.95 | 0.99 | 0.84 | 0.99 |
| Nutlin-3a | 2.0000 | 0.97 | 0.95 | 0.60 | 0.48 | 0.98 | 0.41 | 0.62 | 0.82 | 0.62 | 0.48 | 0.75 | 0.95 | 0.62 | 0.93 |
| Nutlin-3a | 10.0000 | 0.90 | 0.96 | 0.15 | 0.20 | 0.96 | 0.08 | 0.12 | 0.31 | 0.78 | 0.09 | 0.28 | 0.85 | 0.29 | 0.58 |
