## Supplementary Table 4 for "*SAMHD1* mutations in mantle cell lymphoma are recurrent and confer *in vitro* resistance to nucleoside analogues"

Significant effects of drugs on cell viability per gene mutation (t-test). Cytarabine and fludarabine efficacy is reduced in *SAMHD1* mutated samples, nutlin-3a efficacy is reduced in *TP53* mutated samples.

| **Drug** | **gene** | **diff** | **p.value** | **p.adj** |
| --- | --- | --- | --- | --- |
| Cytarabine | *SAMHD1* | 0,203346991 | 2,370314e-06 | 9,481254e-05 |
| Nutlin-3a | *TP53* | 0,155953905 | 1,667259e-04 | 3,334519e-03 |
| Fludarabine | *SAMHD1* | 0,245735306 | 4,014967e-04 | 4,040329e-03 |
| Nutlin-3a | *ATM* | -0,149892351 | 4,040329e-04 | 4,040329e-03 |
| Cytarabine | *NFKBIE* | -0,168438651 | 1,565215e-02 | 1,070363e-01 |
| Cytarabine | *ATM* | -0,116175874 | 9,849718e-02 | 4,377652e-01 |
| Doxorubicine | *NFKBIE* | -0,100830423 | 9,795572e-02 | 4,377652e-01 |
| Fludarabine | *NFKBIE* | -0,170196557 | 1,125949e-01 | 4,409425e-01 |
| Nutlin-3a | *SAMHD1* | 0,065376738 | 2,006707e-01 | 5,733448e-01 |
| Doxorubicine | *KMT2D* | -0,033224610 | 2,348147e-01 | 6,053122e-01 |
| Fludarabine | *ATM* | -0,061345101 | 2,936135e-01 | 6,908552e-01 |
| Doxorubicine | *SAMHD1* | 0,045808245 | 4,013092e-01 | 8,097875e-01 |
| Nutlin-3a | *NFKBIE* | -0,077935830 | 4,108448e-01 | 8,097875e-01 |
| Cytarabine | *TP53* | -0,032851058 | 5,995939e-01 | 8,887010e-01 |
| Doxorubicine | *ATM* | -0,016560338 | 6,574611e-01 | 8,887010e-01 |
| Fludarabine | *KMT2D* | 0,043587434 | 6,148309e-01 | 8,887010e-01 |
| Nutlin-3a | *KMT2D* | -0,016051075 | 8,512887e-01 | 9,533183e-01 |
| Cytarabine | *KMT2D* | -0,009497615 | 9,359900e-01 | 9,854950e-01 |
| Doxorubicine | *TP53* | 0,001548403 | 9,783114e-01 | 9,932908e-01 |
| Fludarabine | *TP53* | 0,000879759 | 9,932908e-01 | 9,932908e-01 |
